# Single-Cell Atlas of Non-Functioning Gonadotroph Tumors and Anterior Pituitary Reveals Lineage Infidelity and Clinically Associated Transcriptional Programs

**DOI:** 10.64898/2026.09.17.752250

**Authors:** Evan Dennis, Gyeong Dae Kim, Yunli Zhou, Clara Alves Pereira, Ye Rim Chang, Gabrielle van der Zee, Anthony Z. Wang, Iris van Mullem, Alla Tsytsykova, Elizabeth Conner, Ethan Tieu, Roy Soberman, Karen K. Miller, Pamela S. Jones, Allegra A. Petti

**Author notes:** **Corresponding Authors:** Allegra A. Petti, PhD, 185 Cambridge Street, CPZN-3800, Boston, MA 02114, Pamela S. Jones, MD. These authors contributed equally (Evan Dennis, Gyeong Dae Kim). Senior authors (Roy Soberman, Karen K. Miller, Pamela S. Jones, Allegra A. Petti).

## Abstract

**Background:** Non-functioning gonadotroph (NFG) tumors are the most common type of non-functioning pituitary adenomas and cause significant morbidity through mass effect. The molecular programs that distinguish NFG tumor cells from normal gonadotrophs and underlie tumor growth remain poorly understood. The aim of this study was to define transcriptional differences between NFG tumors and the normal anterior pituitary gland (APG) at single-cell resolution and to identify programs associated with tumor growth.

**Methods:** We performed single-cell/nucleus RNA-sequencing of 21 NFG tumors and 8 APG samples, followed by integrative analyses of differential gene expression, regulatory networks, and cell–cell communication.

**Results:** We profiled 77,342 APG cells and 152,649 NFG tumor cells. Within the APG, we identified novel marker genes for each neuroendocrine cell type and distinct transcriptional signatures of anterior and posterior pituitary stem cell populations. Compared with normal gonadotrophs, tumor cells shared a core transcriptomic signature conserved across samples despite inter-tumor heterogeneity. Many conserved genes were markers of non-gonadotroph neuroendocrine and stem cells, suggesting that lineage infidelity is a recurrent feature of NFGs. Pseudobulk profiling identified genes positively correlated with tumor volume, including the known oncogenes *CAD*, *BRF2*, and *SOX12*. Finally, tumors showed increased proportions of myeloid, endothelial, and mural cells and crosstalk between tumor, immune, and stromal populations.

**Conclusions:** This study defines transcriptomic signatures of cell identity in the normal APG and NFG tumors. Our findings reveal programs associated with NFG lineage infidelity and tumor growth, highlighting candidate predictors of patient outcomes and potential therapeutic targets.

**Key Points:**

- Single-cell analysis characterizes transcriptomic landscape of APG and NFG tumors.
- Differential expression analysis identifies lineage infidelity in NFG tumors.
- Transcriptomic profiling identifies genes positively correlated with tumor volume.

**Importance of the Study:** This study sequenced more than 152,000 transcriptomes from 21 nonfunctioning gonadotroph (NFG) tumors and 77,000 transcriptomes from the normal adult human anterior pituitary gland (APG). Differential expression analysis of APG cells identified novel marker genes for each neuroendocrine cell type and defined distinct transcriptional signatures of anterior and posterior pituitary stem cells. Comparing these APG cell-type markers with genes differentially expressed in tumor cells revealed that NFGs display transcriptomic characteristics of non-gonadotroph neuroendocrine cells. Pseudobulk profiling identified *FSHB*-high and *FSHB*-low tumors with distinct transcriptional programs, as well as genes positively correlated with tumor volume. We also characterized the NFG tumor microenvironment. Overall, this study provides novel insight into NFG biology, while markers of lineage infidelity and tumor growth may serve as candidate predictors of patient outcomes and potential therapeutic targets. Importantly, these data provide an essential reference dataset for future studies of NFGs and the APG.

## Introduction

Pituitary neuroendocrine tumors (PitNETs) are the most prevalent intracranial neoplasm, comprising 10-20% of intracranial tumors and occurring in up to 20% of individuals^1^. While most PitNETs are relatively benign, many recur, often repeatedly and unpredictably. The most aggressive tumors cause considerable morbidity due to mass effect, resulting in cranial nerve compression syndromes including visual loss, severe headaches, seizures and/or the development of hypopituitarism^2^.

PitNETs are a diverse class of tumors that develop from the anterior pituitary gland (APG), which is comprised of five main cell types, each of which produces a specific hormone. These cell types arise from three developmental lineages, each regulated by a specific transcription factor (TF). Lactotrophs, thyrotrophs, and somatotrophs express the PIT1 TF (encoded by *POU1F1*), while corticotrophs and gonadotrophs express TPIT (encoded by *TBX19*) and SF1 (encoded by *NR5A1*), respectively. Each APG cell type is associated with a corresponding PitNET. In some cases the tumor secretes the cognate hormone, resulting in an endocrine excess syndrome. In others, the tumor is non-functioning, or “silent,” and does not cause an endocrine phenotype, but can cause symptoms due to mass effect. The vast majority of SF-1 positive tumors are non-functioning.

Recent studies have described and classified PitNETs using multi-omic approaches that have provided insight into the molecular heterogeneity underlying these tumors, describing a diversity of cell-types, methylation patterns, and predictive markers of recurrence^3–6^. In this study, we focus on nonfunctioning gonadotroph tumors (NFGs) of the SF1 lineage, the most common type of non-functioning pituitary adenomas^2^. Despite the prevalence of NFGs, the molecular programs underlying NFG tumor growth and their relationship to the normal anterior pituitary gland (APG) are poorly understood. Thus, we conducted sc/snRNA-seq of 21 NFG tumor and 8 APG samples to characterize the transcriptional landscape of NFGs in comparison with the normal pituitary gland.

## Materials and Methods

### Patient Selection and Clinical Variables

The study cohort consisted of 21 patients with NFGs that demonstrated SF1 immunoreactivity by immunohistochemistry. All patients underwent surgical resection of their tumors at Massachusetts General Hospital and provided written informed consent for study participation between 2023 and 2025. Tumors were defined as a true recurrence if there was development of a new tumor mass following complete prior resection. Tumors were defined as invasive if there was radiographic or intraoperative evidence of invasion beyond the immediate anatomical boundaries of the sella into the sphenoid sinus, cavernous sinus, and/or intracranial space. Tumors were manually segmented from MRI scans using 3D Slicer, and tumor volume was calculated using the Segment Statistics tool.

### Single-cell RNA and single-nucleus RNA sequencing sample and library preparation

Detailed sample and library preparation methods are presented in the Supplementary Materials. Briefly, fresh pituitary tumor tissues were collected from the Neurosurgery and Pathology Departments at Massachusetts General Hospital while Frozen normal pituitary tissues were received from the NIH NeuroBioBank at the University of Miami/University of Maryland Brain and Tissue Bank (IRB 2019P002848). After tissue dissociation, library construction, and DNA purification and quantification, libraries were sequenced at Novogene (Sacramento, CA) on an Illumina NextSeq 2000 sequencer using the P3 Flow Cell to achieve a minimum of 20,000 paired-end reads per cell.

### Single-cell and single-nucleus RNA sequencing data analysis

Detailed transcriptomic processing methods are presented in the Supplementary Materials. Briefly, reads were aligned to the Chromium human transcriptome probe set v1.0.1 (GRCh38-2020-A). For each sample, doublets were detected using DoubletFinder (v2.0.4) and removed^7^. Ambient RNA and empty droplets were removed using CellBender^8^. Downstream transcriptomic data processing was performed using Seurat (v5.4.0)^9^. Global filtering was applied by removing cells expressing fewer than 500 unique features, greater than 10% mitochondrial reads, and fewer than 500 or greater than 30,000 UMI’s. Samples were integrated with Harmony via the IntegrateLayers command with method = ‘HarmonyIntegration’^10^. Cells were clustered using Seurat’s pipeline and annotated based on canonical markers. Major cell populations were then subclustered at a higher resolution to identify low-quality clusters before the whole dataset was re-processed. During subclustering of NFG tumor cells, one low-quality cluster with a low feature count was excluded despite meeting the predefined quality-control thresholds.

### Identification of Novel Neuroendocrine Markers

Novel neuroendocrine markers were identified by first running the Seurat command FindAllMarkers on cell types of the APG dataset with a min.pct set to 0.1. Known neuroendocrine markers as defined by manual search, inclusion in the Human Protein Atlas term HPA:0370000 (Pituitary gland), and identification by Cheung et al^11^ were then excluded before visualization.

### Transcription Factor Regulatory Network Analysis

Transcription factor regulatory networks were inferred using the pipeline from the hdWGCNA package (v0.4.08)^12^. Methods are described in greater detail in the Supplementary Materials. TF binding motifs within gene promoter regions were identified using the MotifScan command and the JASPAR 2020 motif database^13^, with additional motifs sourced from the HOCOMOCO database^14^.

### Cell-cell Communication Analysis

Cell-cell communication analysis was conducted using the CellChat package (v2.2.0)^15^. Cell chat objects for normal APG and tumor datasets were created using the create CellChat command with cells grouped by cell type. Ligand-receptor interactions were inferred using the standard CellChat pipeline.

### Identification of Conserved NFG Genes

Upregulated NFG genes conserved across samples were identified by running FindMarkers between tumor cells from each individual NFG sample and gonadotroph cells from the single-cell-sequenced APG sample only with min.pct set to 0.1, Conserved genes were defined as genes with an avg_log2FC > 1 or avg_log2FC < -1 in at least 18 samples and expressed in < 30% of gonadotrophs for upregulated genes and < 30% of cell from each tumor for downregulated genes. Genes were then ranked by mean combined specificity score calculated as pct.1∗(1-pct.2)∗avg_log2FC for each sample for upregulated genes and pct.2∗(1-pct.1)∗avg_log2FC for downregulated genes. Mitochondrial, hemoglobin, and canonical transcription factor/hormone genes were excluded from analysis.

### Ontological and Cell-Type Enrichment

Ontological enrichment analyses were conducted on gene sets throughout the manuscript using the gost command from the gprofiler2 package v(0.2.3)^16^, EnrichR^17^ and MsigDB^18^. Enrichment of SF1 tumor differentially expressed genes (DEGs) within normal neuroendocrine cell types was performed using a hypergeometric test from the hypeR package (v2.2.0)^19^. Detailed methods are included in the Supplemental Materials.

### Pseudobulk Volume Correlation, Differential Expression, and PCA Clustering

Detailed pseudobulked methods are presented in the Supplementary Materials. Briefly, pseudobulked expression profiles from the raw counts of each sample using Seurat’s AggregateExpression() function. Only genes detected in at least 5% of cells in at least five samples were retained. A DESeq2^20^ object was constructed and pseudobulk counts were normalized. Spearman correlations were calculated between transformed gene expression and tumor volume for each gene. Differential expression results for the variable of interest were extracted using a design formula including sex and the variable of interest. For clustering, PCA was run on the top 2,000 variable genes in a pseudobulked DGEList object created with edgeR^21^. Sample-level clustering was performed using k-means clustering with three clusters.

### Differential Expression of Recurrent and Invasive Tumors and Gene Set Activity Scoring

Differential expression was assessed using Seurat’s FindMarkers() function, with min.pct set to 0.20 and min.diff.pct set to 0.10. Genes with an absolute average log₂ fold change > 0.75 and an adjusted P value < 0.01 were then evaluated at the sample level by comparing mean expression between conditions using the Wilcoxon rank-sum test. Genes meeting the significance criteria in both analyses were considered differentially expressed. Differentiation, invasive, and consensus myeloid gene set scores were calculated using the AddModuleScore() function.

### Differential Cell-Type Proportion Analysis

Cell-type proportions between for each sample was assessed by calculating the proportion of cells assigned to each cell type relative to the total number of cells in that sample. Only proportions consisting of greater than 50 cells were included. Differential cell-type proportions between samples in each condition were assessed using a quasibinomial generalized linear model, modeling the number of cells of each cell type relative to all other cells as a function of tumor status. P values were adjusted using false discovery rate, and proportions with an adjusted P value <0.05 were considered significant.

## Results

### Cellular landscape of the anterior pituitary gland and nonfunctioning gonadotroph pituitary neuroendocrine tumors

To characterize the normal APG and NFG tumors, we performed single-cell or single-nucleus RNA sequencing (sc/snRNA-seq) on eight normal APG samples (seven snRNA-seq, one scRNA-seq) and 21 NFG-PitNETs (all scRNA-seq) obtained through transsphenoidal resection (Fig. 1A; Tables S1, S2). Of these tumors, 5 were recurrent and 7 were invasive. Tumors that were either recurrent or invasive were termed aggressive.

**Figure 1.**
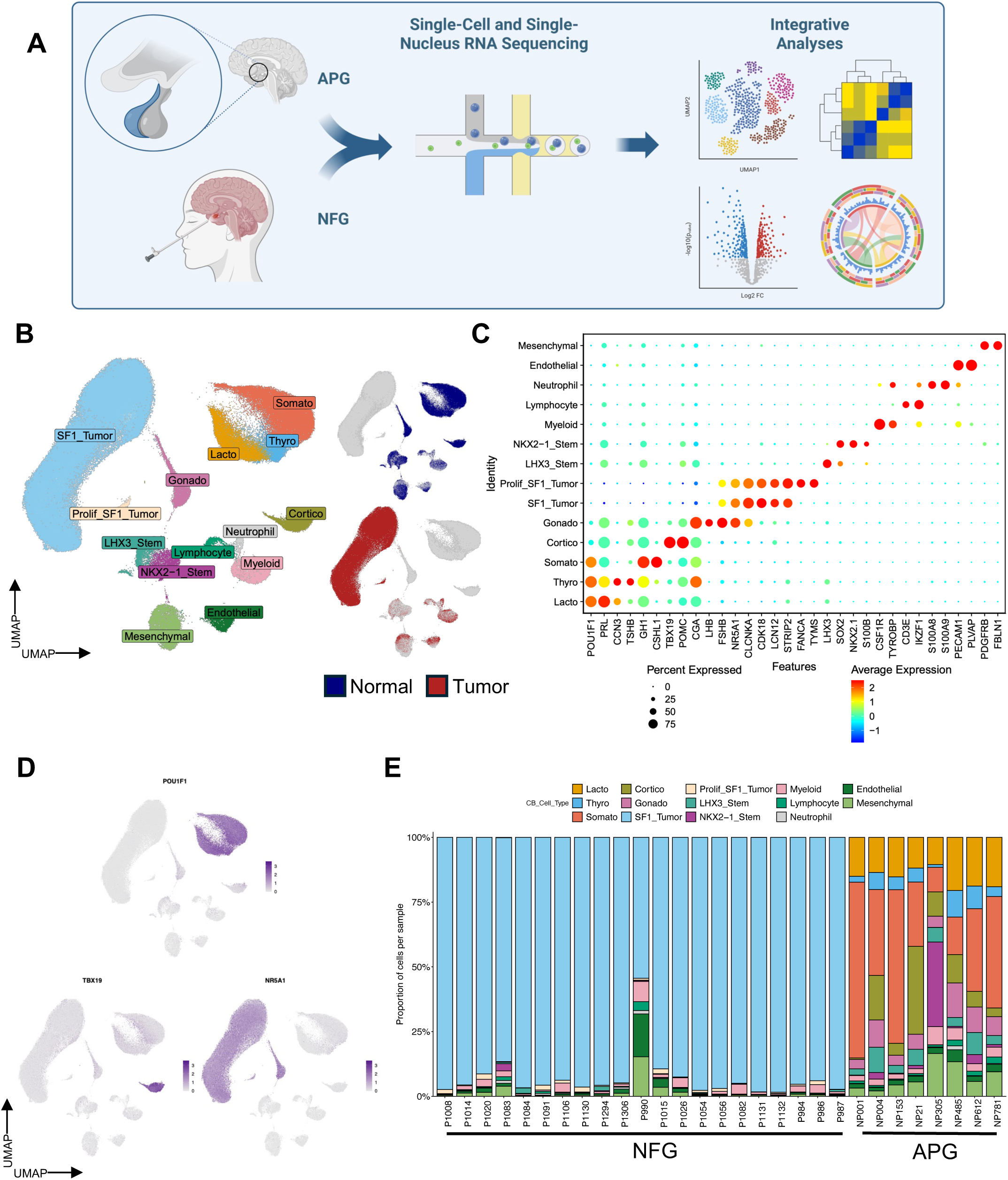
Construction of an integrated single-cell RNA seq dataset of the adult pituitary gland and non-functioning gonadotrophs. **A**. Schematic overview of the study workflow. **B.** Uniform Manifold Approximation and Projection (UMAP) of the integrated tumor and normal dataset. Normal and tumor cells are shown in navy and red, respectively. **C.** Dot Plot showing expression of canonical marker genes across cell types in the integrated tumor/normal dataset. Dot color and size indicate average expression and the percentage of expressing cells, respectively. **D.** Feature Plots of lineage-defining neuroendocrine transcription factors: POU1F1 (PIT-1, top), TBX19 (TPIT, bottom left), NR5A1 (SF1, bottom right). **E.** Bar plot showing distribution of cell types per sample. Panel A Created in BioRender. Tieu, E. (2026) https://BioRender.com/n1y6402 **Alt text:** Workflow schematic and integrated single-cell maps of normal pituitary and tumor samples. Tumor cells form a distinct SF1-expressing cluster separate from normal endocrine, stem, and microenvironmental cells, and tumor samples consist almost entirely of tumor cells.

Following stringent quality control, we retained 77,342 cells from normal APG samples and 152,649 cells from NFG-PitNET samples (Fig. S1A-B). Based on the expression of lineage-specific transcription factors (TFs) and canonical marker genes, we identified all five major neuroendocrine cell types including gonadotrophs expressing *NR5A1* (SF1), corticotrophs expressing *TBX19* (TPIT), somatotrophs expressing *POU1F1* (PIT1) and *GH1*, thyrotrophs expressing *POU1F1* and *TSHB*, and lactotrophs expressing *POU1F1* and *PRL* (Fig. 1B-E). Furthermore, we identified microenvironmental populations across both normal APG and tumor samples, including myeloid cells, neutrophils, lymphocytes, mesenchymal cells, and endothelial cells, along with two distinct stem cell populations, characterized by *LHX3* or *NKX2-1* expression (Fig. 1B).

### Novel marker genes and transcriptional regulatory programs define neuroendocrine cell types

We first focused on the 77,342 APG cells, which comprised the three major endocrine lineages (SF1, TPIT, and PIT1), distinct LHX3⁺ and NKX2-1⁺ stem cell populations, and microenvironmental populations (Fig. 2A-B).

**Figure 2.**
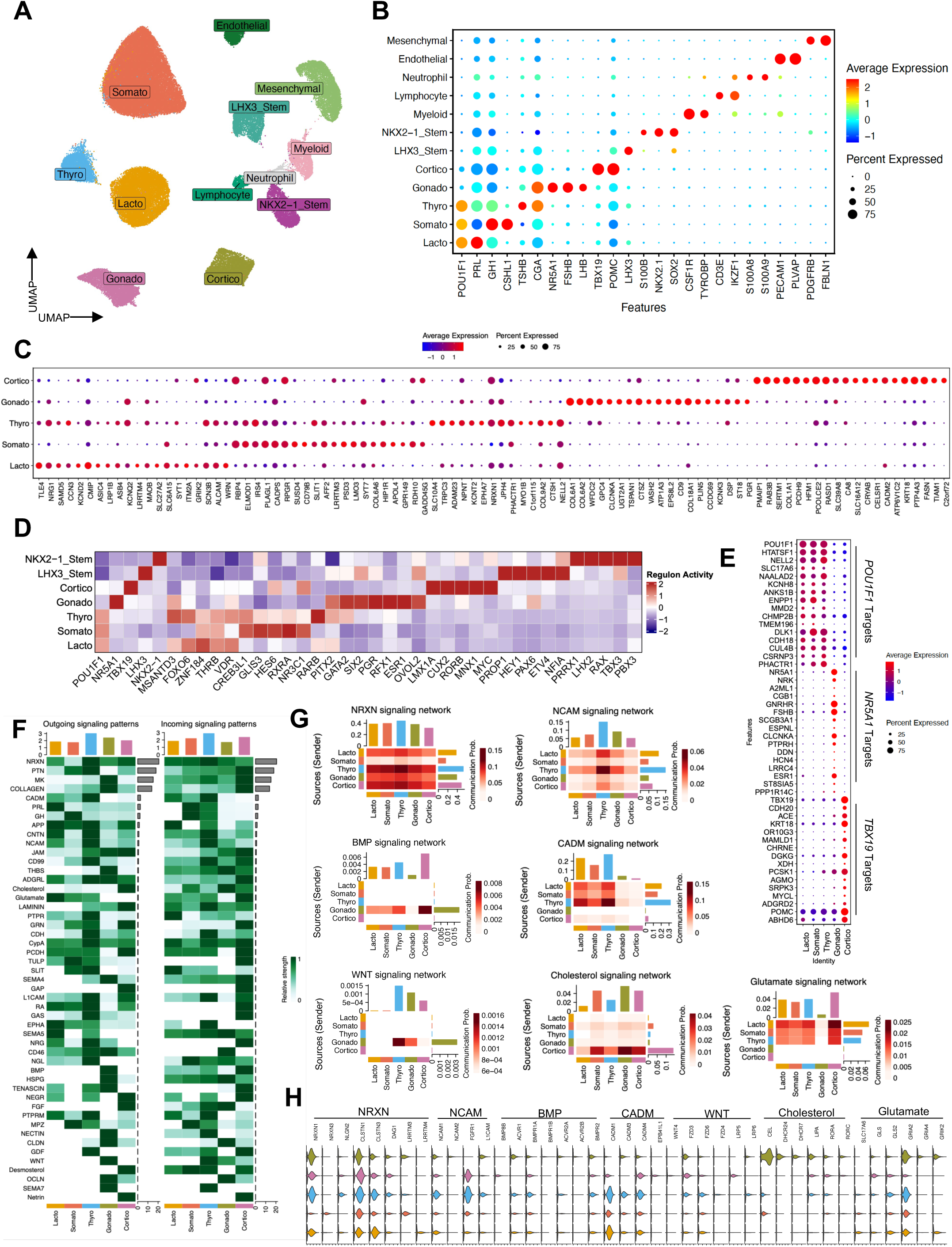
Atlas of the normal pituitary gland expands the transcriptomic profile of neuroendocrine cells. **A.** UMAP of the normal human pituitary gland dataset. **B.** Dot Plot showing expression of canonical marker genes across normal human pituitary cell types. Dot color and size indicate average expression and the percentage of expressing cells, respectively. **C.** Dot Plot showing expression of novel marker genes across five neuroendocrine cell types. **D.** Heatmap of top regulon (transcription factor and its target genes) activity across five neuroendocrine cell types and two stem cell populations. The first five columns show canonical lineage transcription factors for each cell type. **E.** Dot plot showing expression of top target genes of the POU1F1, NR5A1, and TBX19 regulons across five neuroendocrine cell types. **F.** Heat maps showing ligand-receptor signaling from each major neuroendocrine cell type to other neuroendocrine cell types at the aggregate pathway level. **G.** Heatmaps indicating communication probability for selected signaling networks (NRXN, NCAM, BMP, CADM, WNT, Cholesterol, and Glutamate) between neuroendocrine cell types. Rows indicate sender and columns indicate receiver cell types; bar plots on the top and right show the summed incoming and outgoing signaling for each cell type, respectively. **H.** Violin plots showing expression of ligand and receptor genes contributing to the signaling networks in (G) across neuroendocrine cell types. **Alt text:** Single-cell maps, marker plots, and signaling heatmaps of the normal pituitary. Each of the five endocrine cell types and two stem cell populations shows distinct marker genes, lineage transcription factor activity, and patterns of cell–cell signaling.

After filtering out known neuroendocrine cell marker genes, we identified several previously unreported markers exhibiting lineage-specific expression patterns (Fig. 2C). Notably, within the PIT1 lineage, thyrotrophs, somatotrophs, and lactotrophs displayed distinct transcriptional signatures, enabling more precise annotation of these closely related populations (Fig. 2C). Lactotrophs (and to a lesser extent thyrotrophs) specifically express genes associated with synaptic transmission (*NRG1, KCND2, ASIC4, KCNQ2, LRRTM4, SYT1, GRIK2, SCN3B, and SLC6A15*) while corticotrophs were enriched for genes associated with regulated exocytosis (*RAB3B*) and synaptic cell adhesion (*CADM2*)^22^. Somatotrophs were enriched for genes involved in calcium-dependent secretory vesicle exocytosis (*CADPS*, *SYT7*) and excitatory synapse organization (*LRRTM3*)^23,24^. We also uncovered cell-type-specific expression patterns of several collagen genes. For example, *COL6A6* was preferentially expressed in somatotrophs, *COL22A1* in thyrotrophs and gonadotrophs, *COL6A1* and *COL6A2* in gonadotrophs, and *COL1A1* in corticotrophs (Fig. 2C). Pathway analysis of top DEGs for each cell type confirmed enrichment of biological pathways associated with the respective endocrine cell identities, (Fig. S2). For example, somatotrophs showed enrichment of prolactin receptor signaling and vitamin A metabolism, consistent with evidence that retinoic acid regulates GH expression and acts synergistically with thyroid and glucocorticoid hormones in pituitary GH1 cells^25^.

To delineate the transcriptional regulatory programs underlying cell-type identity, we inferred regulons, defined as TFs and their co-expressed target genes, active within each APG population. As expected, regulons associated with canonical lineage-defining TFs (e.g. *POU1F1*, *NR5A1*, *TBX19*) exhibited the highest activity in their corresponding cell types (Fig. 2D). Highlighting the biological fidelity of the inferred regulons, the *NR5A1* regulon included known transcriptional targets of SF1 such as *FSHB* and *GNRHR*, while the *TBX19* included *POMC* along with *PCSK1*, the principal enzyme that cleaves POMC into ACTH (Fig. 2E). We identified additional TFs with established cell-type-specific regulatory activity, providing further insight into the molecular characteristics of each lineage. For example, gonadotrophs showed higher inferred activity of the *GATA2* regulon, which is involved in gonadotroph and thyrotroph specification.

We next characterized the two stem cell populations defined by expression of *NKX2-1* and *LHX3* respectively. Prior work has shown that *LHX3*-expressing stem cells are derived from Rathke’s pouch and ultimately pattern the anterior pituitary gland^26^. Conversely, *NKX2-1* is expressed in the neural ectoderm of ventral diencephalon, which gives rise to the infundibulum^27^. These stem cell populations shared stemness-related pathways, including WNT signaling and morphogenesis, and expressed established stem cell markers such as *SOX2*, *SOX9*, and *CDH2* (Figs. S3A-B). Nevertheless, many genes were differentially expressed between the two populations (Fig. S3C). Consistent with a neural ectodermal origin, NKX2-1^+^ stem cells were enriched for terms related to neurodevelopment, driven by genes such as *LHX2, MAP1B,* and *FAT3*. Conversely, LHX3^+^ stem cells were enriched for pituitary and endocrine developmental programs, driven by increased expression of *LHX3, SIX1,* and *PITX2* (Figs. S3D–E). Regulon-based analysis further underscored these differences, revealing distinct target gene networks for *LHX3* and *NKX2-1* and differential regulon activity between the two populations of other TFs (Fig. S3F, Fig. 2D).

We next used cell-cell interaction analysis to infer ligand-receptor interactions that facilitate communication among these diverse cell types (Table S3). This revealed a mixture of shared and cell-type-specific signaling pathways (Fig. 2F-H). For instance, neuronal adhesion-related interactions mediated by NRXN and NCAM were broadly shared across the neuroendocrine cell types, though the expression of specific ligands and receptors within these pathways varied across cell types (Fig. 2G-H). Conversely, relationships such as BMP, CADM, and WNT interactions showed lineage-specific patterns (Fig. 2G-H). Notably, PRL signals from lactotrophs and GH signals from somatotrophs were inferred to be received by somatotrophs and thyrotrophs (Fig. 2F). Metabolic crosstalk similarly demonstrated cell-type-specific patterns as corticotrophs communicated with all other neuroendocrine cell types via cholesterol signaling, while glutamate signaling was broadly enhanced across the PIT1 lineage, but was received most strongly by corticotrophs. Within the stem cell compartment, LHX3^+^ stem cells uniquely engaged IGFBP, PDGF, and BMP interactions, whereas NKX2-1^+^ stem cells uniquely engaged FGF, EPHA, and SLIT interactions (Fig. S3G). These stem cell populations also demonstrated cross-talk with one another via IGFBP, BMP, FGF, and NOTCH pathways among others (Fig. S3G). Together, these findings suggest that, although these populations share common roles in stem cell maintenance and tissue organization, they exhibit distinct developmental and functional programs.

### Cellular composition and lineage-associated transcriptional programs in NFG tumors

We next independently clustered the 152,649 NFG tumor cells (Fig. 3A). All 21 tumors (8 male and 13 female patients) stained for SF1, but not TPIT or PIT1 (Fig. 3B). *NR5A1* transcripts were detected in all samples at varying levels, whereas *FSHB* showed a dichotomous pattern, with more than half of the tumors expressing substantial *FSHB* despite clinically nonfunctioning status. A subset of tumors weakly expressed alternative lineage markers, such as *TBX19* and *POMC* (P1083) or *PRL* (P1294). Aggressiveness, recurrence, and invasiveness showed no clear association with Ki-67, tumor volume, or expression of canonical lineage-defining transcription factors (Fig. 3B).

**Figure 3.**
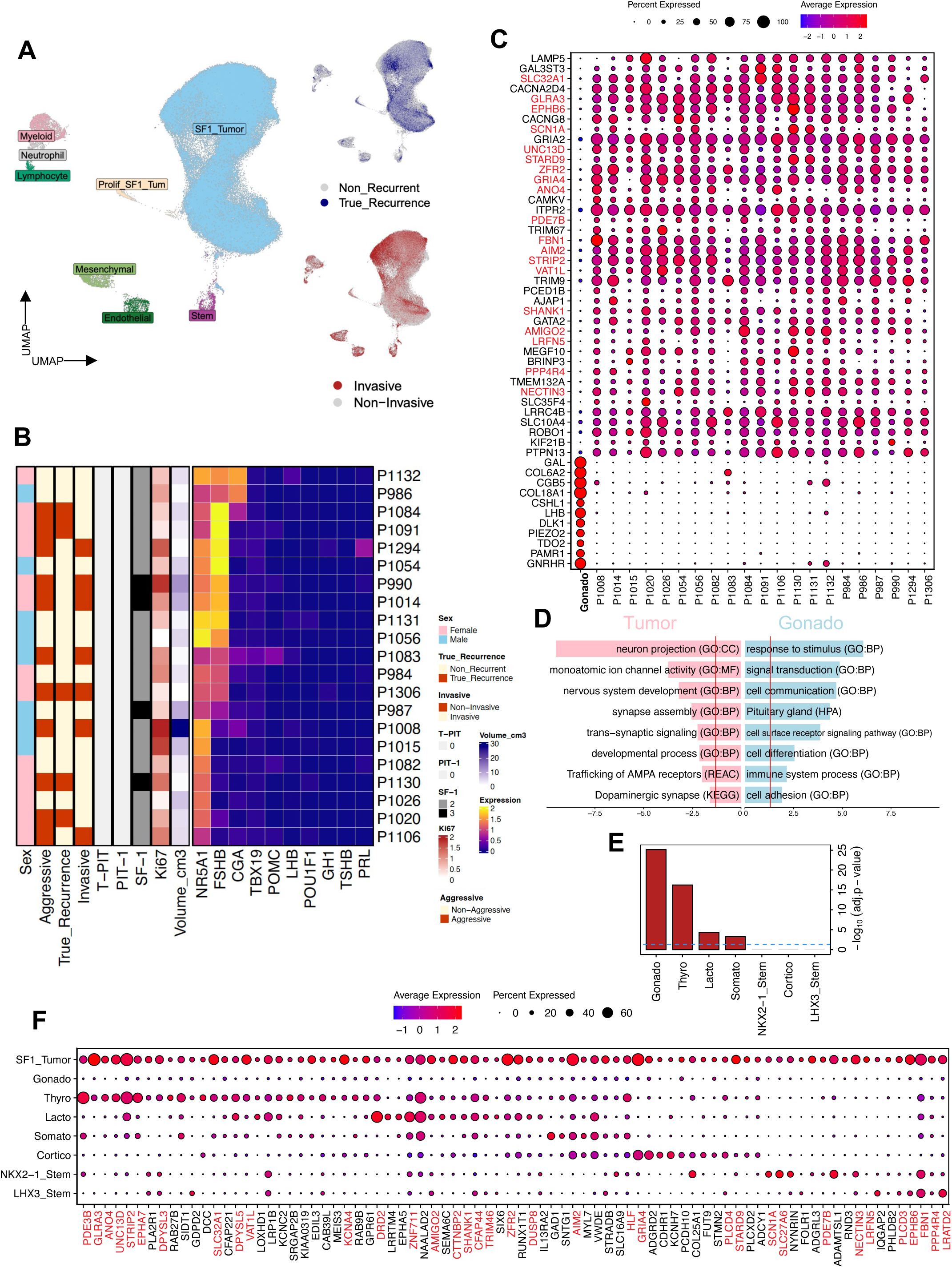
Non-functioning gonadotroph tumors share and diverge from the transcriptional programs of normal neuroendocrine cells. **A.** UMAP of the non-functioning gonadotroph tumor (NFG) dataset. recurrent and invasive cells are shown in navy and red, respectively. **B.** Heatmap showing clinical features (aggressive, recurrence, invasive, and tumor volume) and marker gene expression level at the protein and transcripts per sample. **C.** Dot plot showing expression of top 40 upregulated and top 11 downregulated genes in NFGs compared to gonadotrophs conserved across samples. Red text indicates gene is a marker of a non-gonadotroph pituitary cell type depicted in E. **D.** Bar plot showing ontological enrichment of conserved upregulated genes in NFGs and normal gonadotrophs. Red vertical line indicates significance. **E.** Bar plot showing hypergeometric enrichment of marker genes of pituitary cell types within NFG markers. **F.** Dot plot showing genes with overlapping expression between NFG cells and non-gonadotroph neuroendocrine and stem cell types in the APG. Red text indicates gene is part of the conserved NFG transcriptomic signature. Only the APG sample sequenced with single-cell RNAseq was used for comparison to tumor samples in C-F. **Alt text:** Tumor cell maps, clinical heatmap, and gene expression plots. All tumors express NR5A1, with variable FSHB. Tumor genes overlap most with gonadotroph markers but also with thyrotroph and other lineage markers, and a conserved neural and ion-channel gene signature appears across tumors.

Because transcription factors are central to the function and classification of PitNETs and their cognate neuroendocrine cell lineages, we performed regulon activity analysis to better understand which regulons were active within the tumor cells (Fig. S4A). Along with *NR5A1*, we observed high activity of a variety of transcription factors involved in neuroendocrine differentiation, such as *PITX1*, *PITX2*, *SIX1*, *GATA2*, and *ATF2* required for GnRH-induced *FSHB* expression in gonadotrophs (Fig. S4B). There was also notable activity of the *PBX1* regulon. *PBX1* is increasingly recognized as an oncogenic “master regulator” reactivated in hormone-driven cancers^28^, and small-molecule PBX1 inhibitors are already in early development for other cancers^29^. A thyrotroph signaling axis comprised of *THRA*, and *THRB* was also active.

Differential expression analysis of genes between samples showed that each NFG tumor is defined by a unique transcriptomic profile, demonstrating remarkable inter-tumor heterogeneity (Fig. S5A, B). However, to identify dysregulated genes conserved across samples, we compared the tumor cells from each sample to gonadotrophs in the normal APG and identified shared transcripts upregulated in a large majority of samples (Fig. 3C, S6) (see Methods). This process revealed a conserved transcriptomic signature of NFG dysregulation consisting of 128 genes total (Fig. S6, Table S4). Many of the top upregulated genes have previously been associated with NFGs or pituitary adenomas more broadly including *LAMP5, GAL3ST3, SLC32A1, CACNA2D4, GLRA3, EPHB6, CACNG8, GRIA2, GRIA4*, *STARD9,* and *AMIGO2* (Fig. 3C), providing independent support for the biological relevance of our findings^30–38^. Pathway analysis revealed robust enrichment for genes involved in neuron projection and development (eg. *LAMP5, TRIM9, SHANK1, SEMA6D, EPHA7*) and monoatomic ion channels including AMPA glutamate receptor subunits *GRIA2* and *GRIA4* and alpha subunits of voltage-gated sodium channels *SCN1A, SCN2A,* and *SCN9A* (Fig. 3C-D and S6). Query of the Drug–Gene Interaction Database^39^ identified several agents reported to interact with the ion channel-related genes, as well as with other upregulated transcripts (Fig. S6, Table S5). Top downregulated genes were enriched for terms related to signal transduction and cellular communication, with downregulation of gonadotroph related genes *GAL, LHB*, and *GNRHR* (Fig. 3C-D).

To further delineate lineage-related transcriptional features of SF1+ tumor cells, we assessed the overlap between SF1+ tumor cell DEGs and marker genes of each normal neuroendocrine and stem cell population. As expected, SF1+ tumor cells showed the strongest enrichment for gonadotroph markers, consistent with their gonadotroph origin (Fig. 3E, S4C-D). Surprisingly however, tumor cells also expressed multiple genes specific for non-gonadotroph neuroendocrine cell types. Of these, 35 were part of the NFG signature conserved across samples, including 19 of the top 40 conserved markers (Fig. 3F). For example, after gonadotrophs, thyrotroph markers were most enriched in tumor cells and included the top conserved NFG genes *GLRA3, SLC32A1, ANO4* and *VAT1L* among others. Notably, tumor cells expressed *DRD2* encoding the dopaminergic D2 receptor canonically expressed in lactotrophs while also expressing *GAD1* and *GRIA4,* which were highly specific for somatotrophs and corticotrophs respectively (Fig. 3F). While there was no significant overall enrichment of markers from either stem cell population within tumor cells, several stem cell markers, including *SCN1A, SLC27A6, NECTIN3, EPHB6*, and *FBN1*, were part of the conserved NFG signature. Together, these findings suggest that NFG tumors exhibit lineage infidelity, with many of the top conserved markers that have previously been associated with pathogenicity expressed in other non-gonadotroph pituitary cell types.

### Transcriptional heterogeneity of NFG tumors and clinically associated transcriptional programs

To further investigate intratumoral transcriptional heterogeneity among NFG tumor cells, we subset and re-clustered this population, identifying six tumor cell states (Fig. 4A-D). Each cluster was comprised of cells from multiple samples, though clusters SF1_Tum_C, SF1_Tum_D, and SF1_Tum_E were primarily derived from one or two dominant samples each (Fig. 4A-B). These dominant samples were primarily recurrent in clusters SF1_Tum_C and SF1_Tum_E, and were primarily invasive in SF1_Tum_D (Fig. S7A-B). Functional enrichment analysis of DEGs in each cluster showed enrichment in terms related to cell motility and migration in SF1_Tum_B while cluster SF1_Tum_C was enriched for terms related to cell adhesion and signaling. In contrast, DEGs from the invasive-dominated SF1_Tum_D cluster were enriched for terms related to cell population proliferation (Fig. 4C).

**Figure 4.**
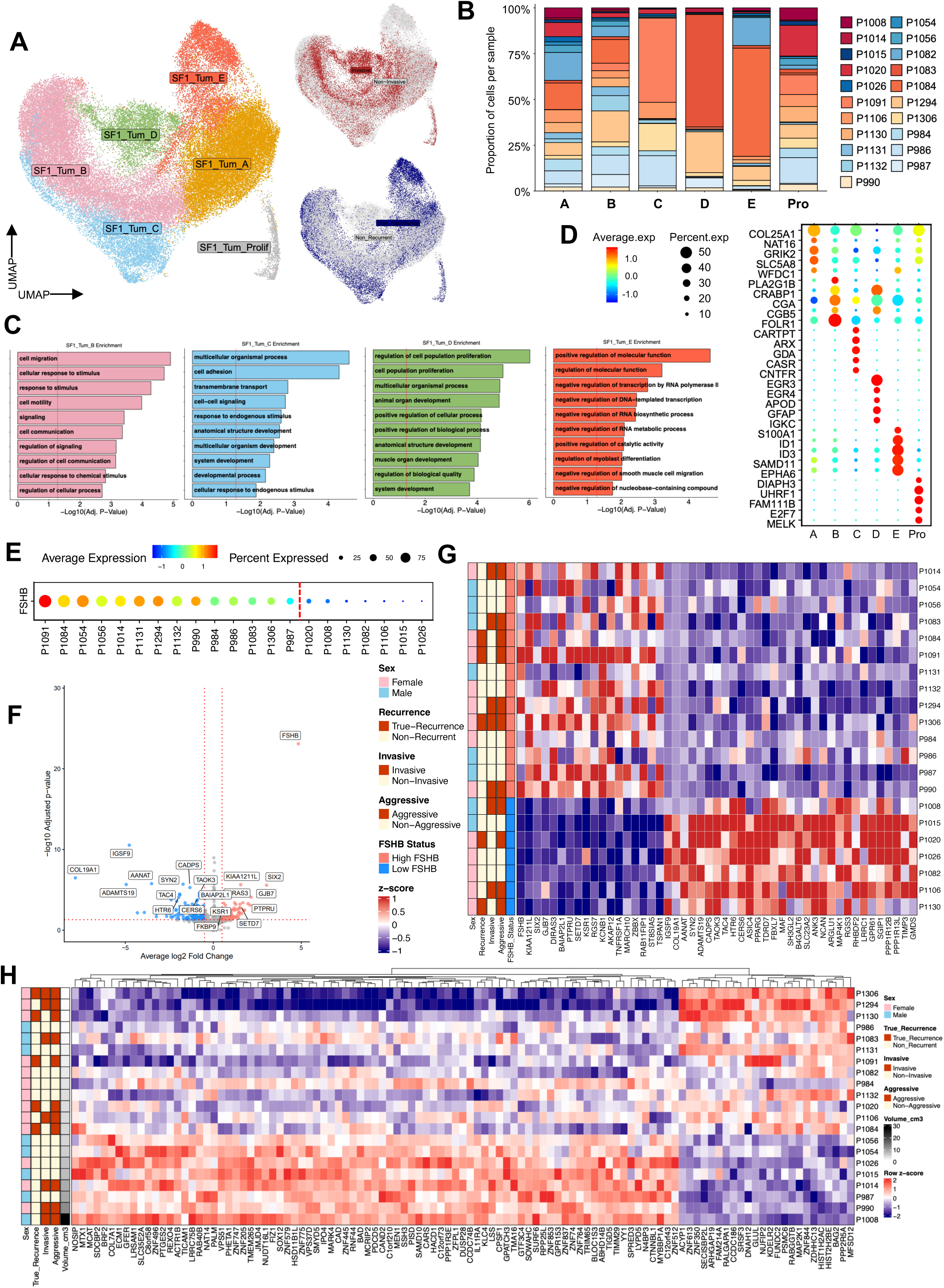
Intratumoral heterogeneity and clinical subgroup-specific characteristics of non-functioning gonadotroph tumor cells. **A.** UMAP of six SF1 tumor cell subclusters. Recurrent and invasive cells are shown in navy and red, respectively. **B.** Stacked bar plot showing the proportion of cells contributed by each sample within each tumor subcluster. **C.** GO enrichment of upregulated genes in subclusters SF1_Tum_B, SF1_Tum_C, SF1_Tum_D, and SF1_Tum_E. Red vertical line indicates significance. **D.** Dot plot showing marker genes of each tumor subcluster. Dot color and size indicate average expression and the percentage of expressing cells, respectively. **E.** Dot plot showing *FSHB* expression in tumor cells per sample. Samples with less than 40% of cells expressing *FSHB* were classified as low_FSHB. **F.** Volcano plot of differentially expressed genes identified by pseudobulk analysis between high_FSHB and low_FSHB tumors. **G.** Heatmap showing scaled normalized pseudobulked counts of the top 50 significantly differentially expressed genes per group identified by pseudobulk analysis between high_FSHB and low_FSHB tumors per sample. **H.** Heatmap showing scaled normalized pseudobulked counts per sample of genes significantly correlated with tumor volume per sample. Row annotations indicate sex, recurrence, invasiveness, aggressiveness, and tumor volume. **Alt text**: Tumor subcluster maps, sample composition bars, enrichment charts, and expression heatmaps. Tumor cells form six subclusters, some dominated by single tumors. Tumors split into FSHB-high and FSHB-low groups with distinct expression, and a set of genes correlates with tumor volume.

Inspection of expression of key genes related to normal pituitary function had demonstrated that many of our NFG samples expressed *FSHB* despite being non-functioning tumors (Fig. 3C, Fig. 4E). However, robust *FSHB* expression was not observed across all samples, with some expressing very low levels of the gene. Pseudobulked differential expression analysis between *FSHB* high and low samples identified 285 significant DEGs, demonstrating distinct transcriptomic profiles defining these two tumor classes (Fig. 4F-G, Fig. S7C, Table S6). *FSHB*-high tumors were enriched for ontological terms related to cell signaling and secretion, with increased expression of chromogranin/secretogranin genes *CHGB, CHGA*, and *SCG2* involved in secretory granule biogenesis (Fig. S7C-D). In contrast, *FSHB*-low tumors were most enriched for terms related to neuron projection and development. These findings suggest that *FSHB* expression may distinguish biologically distinct states within SF1 tumors.

Beyond *FSHB* expression, per-sample pseudobulked expression profiles of multiple genes were significantly positively or negatively correlated with tumor volume (Fig 4H). Notably, *BRF2* and *SOX12* were positively correlated with tumor volume. Both genes have been reported to promote metastasis and invasion in other cancers, through activation of Wnt/β-catenin signaling and enhancement of regulatory T-cell infiltration, respectively^40,41^. In addition, *CAD*, a known cancer related gene that is activated downstream of mTORC1 to support cell growth^42^ along with 16 zinc finger transcription factors were also associated with tumor volume. In contrast, *RALGAPA1*, a negative regulator of Ral GTPases that counteracts oncogenic Ras signaling^43^, was associated with lower tumor volume.

Pseudobulk differential expression analysis between recurrent and non-recurrent samples identified 8 dysregulated genes in recurrent tumors (Fig. S8A-B). Notably, we observed a downregulation of *SHH* in recurrent tumors, consistent with prior literature suggesting that SHH inhibits cell proliferation in pituitary adenomas^44^, though *SHH* had relatively low expression overall. Recurrent tumors also demonstrated upregulation of *EDA*, a member of the tumor necrosis factor family of ligands suggested to play a role in tumor pathogenesis^45^. As pseudobulk analysis is a conservative method, we also identified genes that were differentially expressed at the single-cell level while also demonstrating significant differences between samples (see Methods). This approach identified 40 genes that differed significantly between recurrent and non-recurrent tumors (Fig. S8C). Recurrent tumors were enriched for pathways related to the regulation of stem-cell pluripotency and bone morphogenetic protein signaling, both of which may contribute to tumor-cell plasticity. For example, *SMAD9* and *FOXO1* are two transcriptional regulators driving enrichment within these pathways. A previously reported “well-differentiated” SF1 tumor signature (described by Zhang et al.^13^) showed elevated, though not quite significant activity in non-recurrent tumors, with two genes, *TGFBR3L* and *CRABP1,* overlapping between the well-differentiated gene list and our expanded non-recurrent signature (Fig. S8D-G).

Following a similar strategy, we analyzed non-invasive and invasive tumors. Pseudobulk analysis identified 10 DEGs including *VIM* (Fig.S9A-B). *VIM* is a classic marker of epithelial-mesenchymal transition and has previously been associated with invasive PitNETs^46^. At the single-cell level, non-invasive tumors were enriched for sphingolipid catabolism and insulin-processing pathways, whereas invasive tumors showed elevated TNF-α signaling via NF-κB, AP-1 transcriptional network activity (Fig. S9C, Table S7). Consistent with this, core members of the AP-1 family, including *JUN*, *FOS*, and *FOSB* were significantly upregulated in invasive tumors (Fig. S9D). *LHX3*, which was highly expressed in the stem cell population, was also elevated in invasive tumors (Fig. S9E). To assess the relationship between invasiveness and stem-like transcriptional states, we calculated gene-set scores for invasive- and non-invasive-associated genes in the two stem cell populations (Fig. S9F). The invasive-associated signature was significantly enriched in both LHX3^+^ and NKX2-1^+^ stem cells.

We next investigated whether there are transcriptionally related subsets of NFGs unrelated to clinical characteristics. We generated pseudobulk profiles of each sample and performed principal component analysis and clustering, which yielded three clusters (Fig. S10A). These clusters did not clearly segregate by invasive or recurrent status, although both tumors in C3 were recurrent (Fig. S10B). Each cluster exhibited a distinct profile (Fig. S10C, Table S8). C2 comprised 15 tumors and was characterized by 151 DEGs, including *TMEM196*, *CX3CL1*, *STOX2*, and *MEF2A* (Fig. S10C). Enrichment analysis highlighted cell population proliferation and regulation of insulin secretion (Fig. S10D). C1 comprised five tumor samples and was characterized by 96 DEGs, including *LHX3* and the corticotroph lineage transcription factor *TBX19* (Fig. S10C) with enrichment of developmental and synaptic terms suggested a prominent developmental and neuronal transcriptional program (Fig. S10D). C3 comprised only two tumors but exhibited 414 uniquely upregulated genes, predominantly located on the X chromosome. Consistent with this finding, enrichment analysis highlighted X-linked inheritance, suggesting altered X-chromosome regulation (Fig. S10D).

### NFG PitNETs exhibit robust immune infiltration and heterotypic interactions with immune, vascular, and mesenchymal cells

To define the composition of NFG-PitNETs and normal APGs beyond neuro-endocrine cell populations, we isolated and clustered non-neuroendocrine cells, identifying immune, stem, and vascular cells of the tumor and APG microenvironment (Fig. 5A, Fig. S11A). Compared with APG samples, tumor samples exhibited increased relative proportions of myeloid populations, neutrophils, NK cells, and B cells, indicating enhanced immune-cell infiltration and remodeling of the tumor immune microenvironment (Fig. 5B). Endothelial and mural cells were also more abundant in tumors while fibroblasts were depleted, suggesting vascular remodeling within the tumor micro-environment (TME).

**Figure 5.**
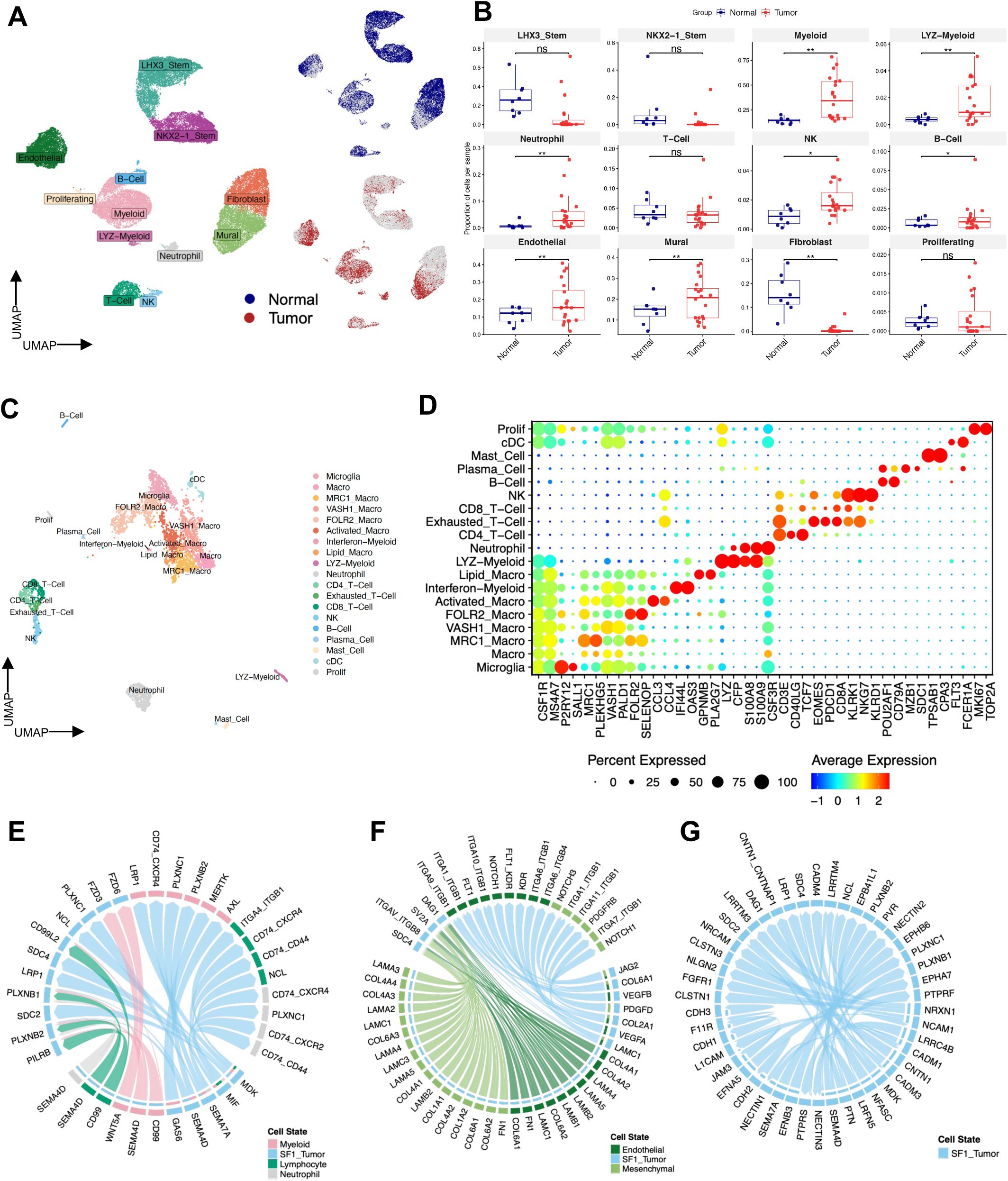
Characterizing NFG-microenvironment interactions and microenvironment cell types. **A.** UMAP of integrated microenvironment cells from NFG and normal adult pituitary gland (APG) samples. Normal and tumor cells are shown in navy and red, respectively. **B**. Box plots showing the proportion of each microenvironment cell type per sample in NFG and normal APG samples. **C.** UMAP of immune cells from NFG samples. **D.** Dot plot showing canonical marker gene expression across various immune cell subtypes. **E-G.** Cell-cell communication between tumor cells and immune cells **(E)**, endothelial and mesenchymal cells **(F)**, and other tumor cells **(G)** at the aggregate pathway level. **Alt text:** Microenvironment cell maps, cell-proportion box plots, and ligand–receptor chord diagrams. Compared with normal pituitary, tumors contain more myeloid, endothelial, and mural cells and fewer fibroblasts, with predicted tumor signaling to immune, vascular, and stromal cells.

We next analyzed the relationship between tumor aggressiveness and TME composition (Fig. S11B-C). At the sample level, aggressive tumors showed a significantly lower relative proportion of myeloid cells than nonaggressive tumors, suggesting that myeloid composition may correlate with the clinical behavior of NFG-PitNETs (Fig. S11D).

To further resolve the tumor-immune compartment, we subclustered tumor-associated immune cells and identified microglia-like cells, multiple myeloid subtypes, and lymphocyte populations (Figs. 5C-D). Myeloid subtypes were distinguished by differential expression of *MRC1*, *VASH2*, and *FOLR2*. We also identified activated myeloid populations expressing the inflammatory chemokines *CCL3* and *CCL4*, as well as smaller populations characterized by interferon-stimulated genes such as *IFI44L* and *OAS3*, the lipid metabolism-associated gene *PLA2G7*, or high *LYZ* expression. Scoring of previously defined myeloid consensus programs^47^ in our myeloid subtypes suggested that the prominent *FOLR2* and *MRC1* myeloid populations possess an immunosuppressive character (Fig. S11E).The lymphoid compartment included CD4 T cells, CD8 T cells, NK cells, and a small population of *PDCD1*-expressing exhausted-like T cells.

We then characterized ligand-receptor interactions between tumor cells and each TME cell population, focusing initially on interactions of tumor cells with lymphocytes, myeloid cells, and neutrophils (Fig. 5E, Table S9). The *MIF*→*CD74*/*CXCR4*/*CD44* axis, associated with macrophage recruitment and polarization, T-cell modulation, and pro-survival signaling, was prominent. Additional immunosuppressive and tumor-supportive interactions included *MDK*→*LRP1* and *GAS6*→*AXL*/*MERTK* signaling. Furthermore, tumor-stromal crosstalk with endothelial and mesenchymal populations was characterized by pro-angiogenic and extracellular matrix (ECM) remodeling pathways such as VEGF (*VEGFA*/*VEGFB*→*FLT1*/*KDR*), *PDGFD*→*PDGFRB,* and collagen and fibronectin (*FN1*) to integrins and syndecans (*ITGAV, ITGB8, SDC4*) (Fig. 5F). Among tumor cells, interactions involving CADM, NCAM, NECTIN, NRXN, JAM, L1CAM pathways and cadherins suggested prominent roles in tumor-cell cohesion whereas *SEMA4*/*7*, and ephrin signaling are associated with spatial organization and migration (Fig. 5G). Together, these findings demonstrate substantial immune cell heterogeneity and complementary cell-cell communication programs within NFG TMEs.

## Discussion

In this study, we profiled 21 NFGs and eight normal APGs at single-cell resolution. Within APGs, we identified cell type-specific markers and regulons, inferred heterotypic interactions, and profiled two distinct stem cell populations. Within NFG PitNETs, we identified a conserved NFG transcriptomic signature, markers of lineage infidelity, transcriptional correlates of clinical properties, and marked perturbations of the microenvironment.

Comparison of NFG tumor cells to gonadotrophs from the APG revealed a transcriptomic signature largely conserved across NFG samples despite marked inter-tumor heterogeneity. Several of the top conserved markers have previously been associated with non-functioning gonadotrophs corroborating the biological relevance of our findings^30–38^. Ontological analysis showed that the conserved signature was dominated by genes involved in neuron projection, nervous system development, and monoatomic ion channel activity, consistent with bulk transcriptomic studies identifying calcium signaling, glutamate receptor, and synapse organization networks as defining features of non-functioning adenomas^33,48^. These programs may be functionally relevant, as spontaneous calcium activity correlates with recurrence of non-functioning adenomas^38^, and in other tumors, co-opted neural programs promote growth and invasion^49^. For instance, AMPA receptor-mediated currents drive glioma proliferation and invasion^50,51^, and aberrantly expressed voltage-gated sodium channels, including Nav1.7 encoded by *SCN9A*, enhance cancer cell migration and invasiveness^52,53^. The conserved upregulation of AMPA receptor subunits (*GRIA2, GRIA4*) and sodium channel α subunits (*SCN1A, SCN2A, SCN9A*) therefore raises the possibility that ion channel activity contributes to NFG growth. Notably, many of these channels are targeted by approved drugs, including the AMPA receptor antagonist perampanel and sodium channel-blocking antiepileptics (Table S5), nominating them as candidates for functional validation.

Remarkably, comparison of NFG DEGs to markers of other pituitary cell types demonstrated that NFG tumors express genes specific for non-gonadotroph neuroendocrine and stem cells. 35 of these lineage-incongruent markers were included in the conserved NFG signature, suggesting lineage infidelity is a recurrent feature of SF1 adenomas. Lineage infidelity in PitNETs has largely been defined by immunohistochemistry, with the WHO 2022 guidelines classifying tumors expressing more than one lineage-defining transcription factor under the “plurihormonal” category^54^. The term “multilineage” has also been proposed with PIT1/SF1 co-expression being the most frequent combination^54^. SF1-positive tumors that express an additional lineage transcription factor are more often invasive, less often gross-totally resected, and more likely to recur than tumors expressing SF1 alone^55,56^. Notably, the lineage-incongruent genes in our conserved NFG signature were not lineage-defining transcription factors, and would therefore not be captured by routine immunohistochemical workup. Our data thus suggest that transcriptional evidence of mixed-lineage differentiation may be present even in tumors classified as SF1-lineage by conventional pathology, and could contribute to their heterogeneity in clinical outcomes.

*FSHB* expression defined an axis of tumor heterogeneity, with some samples expressing robust levels of *FSHB* despite being functionally inactive. In contrast, there was almost no expression of *LHB*. These results are consistent with those of Katznelson et al., who detected *FSHB* expression in 49% of clinically nonfunctioning adenomas using Northern blot analysis, compared with *LHB* expression in only 1% of samples^57^. Additionally, *FSHB* status was not correlated with tumor size or invasiveness, again consistent with prior studies^58^. However, several genes were differentially expressed between *FSHB*-high and *FSHB*-low groups suggesting that *FSHB* expression defines molecularly distinct tumor subtypes. *FSHB*-high tumors were enriched for secretory and signaling programs, whereas *FSHB*-low tumors were enriched for neuronal programs, potentially reflecting different degrees of gonadotroph differentiation and paralleling the differentiation-based subtypes described across PitNET lineages^4^.

Additionally, tumor volume showed a strong relationship with sample-level transcriptional variation, with 83 genes positively correlating with tumor volume. Notably, *CAD*, *BRF2* and *SOX12* were all positively correlated with tumor volume and have previously been shown to be involved in tumor growth^40–42^. Additionally, 16 zinc-finger transcription factors were correlated with tumor size, reflecting the roles of zinc finer proteins related to cancer progression in other tumor types^59^. Conversely, *RALGAPA1*, which negatively regulates RALA and RALB signaling was negatively correlated with tumor volume^43^, raising the possibility that reduced restraint of Ral signaling accompanies larger tumors. Because morbidity in NFG-PitNETs is driven by mass effect rather than hormone excess, these volume-associated programs are directly relevant to the clinical consequences of these tumors and warrant validation as candidate markers of tumor growth.

Limitations of this study include a relatively small cohort for clinical subgroup analysis, the donor dominance of some stem and tumor-cell states, and the absence of an independent validation cohort. Within these limits, this study establishes a single-cell atlas linking normal pituitary lineage identity to the molecular heterogeneity and clinical behavior of NFG-PitNETs. Our findings support a model in which NFGs express a core dysregulation program while acquiring variable lineage infidelity. We also define axes of heterogeneity related to tumor volume and *FSHB* expression. Collectively, these findings provide a foundation for developing molecular classifiers and prioritizing candidate biomarkers and therapeutic vulnerabilities in NFG-PitNETs.

## Ethics

All patients provided written informed consent for study participation. Fresh pituitary tumor tissues were collected from the Neurosurgery and Pathology Departments at Massachusetts General Hospital, approved by the Mass General Brigham Institutional Review Board (IRB 2019P002925). Analysis of these samples and associated clinical data was conducted under a separate IRB-approved secondary-use protocol (protocol #2024P003599). Frozen normal pituitary tissues were received from the NIH NeuroBioBank at the University of Miami/University of Maryland Brain and Tissue Bank (IRB 2019P002848). The use of these samples for gene expression analysis was approved by the IRB (2024P003599).

## Author Contributions

Conceptualization: KKM, PSJ, AAP. Data Curation: PSJ, YZ, YRC, ED, GDK, CAP, IM, EC, AAP. Formal Analysis: ED, GDK, CAP, GZ. Funding Acquisition: KKM, RS, PSJ. Investigation: ED, GDK, YZ, CAP, GZ, IM, AT, YRC, PSJ. Methodology: YZ, ED, IM, AAP. Project Administration: KKM, RS. Resources: PSJ, EC, ET. Software: ED, GDK, GZ. Supervision: KKM, PSJ, AAP. Writing, first draft: GDK, ED, AAP. Writing, review and editing: KKM, PSJ, YRC, YZ.

## Data Availability

Upon publication, processed data will be available on Zenodo (DOI 10.5281/zenodo.18790584) and raw data will be available on GEO. Code will be deposited in Github.

## Acknowledgments

We gratefully acknowledge Fred Ringler and Pamela McLaughlin for their support.

## Declaration of generative AI and AI-assisted technologies use

During the preparation of this work, the authors used artificial intelligence (AI)-assisted tools to adjust code for graph generation and to improve the readability, language, and grammar of the text. After using these tools, the authors reviewed and edited the content as needed and take full responsibility for the content of the publication.

