## Supplemental Figures and Legend for "Single-Cell Atlas of Non-Functioning Gonadotroph Tumors and Anterior Pituitary Reveals Lineage Infidelity and Clinically Associated Transcriptional Programs"

**Single-Cell Analysis of Non-Functioning Gonadotroph Tumors Identifies Lineage Infidelity and Tumor Growth Programs**

**Author Notes**

*These authors contributed equally

**Senior authors

+Corresponding Authors

**Supplementary Methods**

**Single-cell RNA and single-nucleus RNA sequencing sample and library preparation**

Fresh pituitary tumor tissues were collected from the Neurosurgery and Pathology Departments at Massachusetts General Hospital, approved by the Mass General Brigham Institutional Review Board (IRB 2019P002925). Frozen normal pituitary tissues were received from the NIH NeuroBioBank at the University of Miami/University of Maryland Brain and Tissue Bank (IRB 2019P002848). Normal pituitary tissues were from donors who died from non-pituitary-related diseases (Supp. Table 2). The use of these samples for gene expression analysis was approved by the IRB (2024P003599). Pituitary tumor tissues were immediately processed upon release from the Pathology. For frozen normal pituitary tissues from NIH NBB, total RNA was first isolated from a portion of each tissue, and RNA integrity number (RIN) was determined using a Bioanalyzer. Only samples with a RIN greater than or equal to 8 were included in the study.

Single-cell and single-nucleus cDNA libraries were generated from pituitary tumors and normal pituitaries, using the 10X Genomics Chromium Fixed RNA Profiling (aka “Flex”) workflow (10X Genomics, Pleasanton, CA). The Flex fixed RNA approach allowed us to include samples after their pathology phenotypes were determined. Briefly, approximately 20-25 mg of fresh tumor tissue or frozen normal pituitary tissue was minced, fixed, and dissociated into single cells or nuclei using the Chromium next GEM single cell fixed RNA sample preparation kit from 10X Genomics (PN-1000414). The scRNA and snRNA 4-plex libraries were constructed using the Chromium fixed RNA human transcriptome kit (4 rxns x 4 BC, PN-1000475) at the Broad Technology Space - Single Cell Platform (Broad Institute, Cambridge, MA). For each library, 3 × 105 cells or nuclei from each of 4 samples were separately hybridized with 4 individual barcoded probes. After the post-hybridization wash, an equal number of cells (nuclei)/barcoded probe from each hybridization reaction was pooled. To recover 10,000 cells per sample, 6.6 x 104 cells from the cell pool were loaded into a Chromium Next GEM Chip Q. GEM were generated using a Chromium X instrument. GEM recovery, incubation, and pre-amplification were performed according to the reagent kit’s instructions. The cDNA was cleaned up using the SPRIselect reagent from Beckman Coulter (Indianapolis, IN). The final library was constructed by amplifying the cDNA using a set of sample index primers from a Dual Index Kit TS plate (10X Genomics, PN-1000251) following the reagent kit’s instructions. The index PCR was set to 9 cycles. After DNA purification and quantification, the library was sequenced (with four multiplexed samples per run) at Novogene (Sacramento, CA) on an Illumina NextSeq 2000 sequencer using the P3 Flow Cell to achieve a minimum of 20,000 paired-end reads per cell.

**Single-cell and single-nucleus RNA sequencing data analysis**

Sequencing data were obtained from Novogene in the fastq format and were demultiplexed and processed with the ‘multi’ pipeline of CellRanger v7.1.0 or v9.0.1. Reads were aligned to the Chromium human transcriptome probe set v1.0.1 (GRCh38-2020-A). For each sample, doublets were detected using DoubletFinder (v2.0.4) and removed.73 Ambient RNA and empty droplets were removed using CellBender.74 Downstream transcriptomic data processing was performed using Seurat (v5.4.0).75 CellBender output for each sample was merged into a Seurat object. Global filtering was applied by removing cells expressing fewer than 500 unique features, greater than 10% mitochondrial reads, and fewer than 500 or greater than 30,000 UMI’s. The count expression values were then normalized with the NormalizeData command. 2500 variable features were identified using FindVariableFeatures and scaled using the ScaleData command. After running PCA, layers of the Seurat object were integrated with Harmony via the IntegrateLayers command with method = ‘HarmonyIntegration’.76 After identifying neighbors and running UMAP on 25 harmonized dimensions using the FindNeighbors and RunUMAP commands, cells were clustered at a resolution of 2 with FindClusters. Clusters were then annotated based on canonical markers. Following annotation, tumor cells, neuroendocrine cells (comprised of gonadotrophs, lactotrophs, somatotrophs, thyrotrophs, and corticotrophs), and microenvironment cells were each subclustered separately at a higher resolution to identify low-quality clusters designated by high expression of aberrant marker genes of other cell classes or low number of unique features and total counts expressed overall. After removal of low-quality cells, the whole dataset was re-clustered using the FindClusters function at a resolution of 2 with resulting clusters once again annotated based on canonical markers. For each subclustered dataset, the selected cell populations were reprocessed as above from the raw count data. During subclustering of NFG tumor cells, one cluster with a low feature count was excluded despite meeting the predefined quality-control thresholds before further subclustering.

**Transcription Factor Regulatory Network Analysis**

Transcription factor regulatory networks were inferred using the standard pipeline from the hdWGCNA package (v0.4.08). Genes expressed in at least 5% of cells in at least one cell type were included. Meta cells were generated with the MetacellsByGroups command from cells grouped by cell type and condition, using a nearest-neighbor parameter k = 25 and a max_shared parameter of 10. Following hdWGCNA analysis, transcription factor analysis was conducted on the resulting hdWGCNA object using the standard pipeline. TF binding motifs within gene promoter regions were identified using the MotifScan command and the JASPAR 2020 motif database,79 with additional motifs sourced from the HOCOMOCO database.80 Regulons were defined by using the AssignTFRegulons command with strategy = “A” (where each gene is assigned to the regulon of its top n TF’s), reg_thresh = 0.01, and n_tfs = 10. Positive and negative regulon scores were then calculated for each cell using the RegulonScores command and differential regulon expression between epithelial and non-epithelial cells was calculated using the FindDifferential Regulons command.

**Ontological and Cell-Type Enrichment**

Ontological enrichment analyses were conducted on gene sets throughout the manuscript using the gost command from the gprofiler2 package v(0.2.3) EnrichR and MsigDB. Enrichment of SF1 tumor differentially expressed genes (DEGs) within normal neuroendocrine cell types was performed using a hypergeometric test from the hypeR package (v2.2.0). DEGs in neuroendocrine cell types were identified in the single-cell APG sample only using the FindAllMarkers command with min.pct of 0.15. Only the single-cell APG sample was used for comparison with tumor markers because all tumor samples were profiled using single-cell, rather than single-nucleus, RNA sequencing. SF1 tumor cell-type genes were defined as genes differentially expressed in the SF1 tumor cluster relative to neuroendocrine and stemm cells from the single-cell APG sample, identified using the FindAllMarkers function min.pct of 0.2. Significant positively differentially expressed genes in each group were then filtered to include genes with a log2 fold-change greater than 1.5 and expression in at least 20% of cells in a population. Genes were then ordered based on a combined score calculated as pct.1∗(1-pct.2)∗avg_log2FC, and the top 200 genes for each cell type were selected. The background number of genes was 20,000. All p-values were corrected for multiple hypotheses using the FDR method.

**Volume Correlation Analysis**

Volume correlation analysis was conducted by first creating pseudobulked expression profiles from the raw counts of each sample using Seurat’s AggregateExpression() function. Only genes detected in at least 5% of cells in at least five samples were retained. A DESeq2 object was constructed and pseudobulk counts were normalized using DESeq2 size-factor estimation and variance-stabilizing transformation.^44^ Spearman correlations were calculated between transformed gene expression and tumor volume for each gene. P values were adjusted using false discovery rate, and genes with an adjusted P value <0.05 were considered significant.

**Pseudobulk Differential Expression Analysis**

Differential expression analysis between different types of NFG tumor samples was conducted by first creating pseudobulked expression profiles from the raw counts of each sample using Seurat’s AggregateExpression() function. Only genes detected in at least 5% of cells in at least five samples were retained. A DESeq2 object was constructed using a design formula including sex and the variable of interest (~ Sex + VOI). Differential expression results for the variable of interest were extracted and Log2 fold-change estimates were subsequently shrunk using the adaptive shrinkage estimator implemented in apeglm. Genes were considered significantly differentially expressed if they had an absolute log2 fold change > 0.5 and an adjusted p value < 0.05. For pseudobulk differential expression between high and low *FSHB* samples, samples with less than 40% of cells expressing *FSHB* were classified as low_FSHB.

**PCA clustering of Pseudobulk Tumor Samples**

Raw single-cell counts were summed across cells within each sample to generate sample-level pseudobulk profiles. Only genes detected in at least 5% of cells in at least five samples were retained. A DGEList object was created and libraries were normalized using TMM normalization with the calcNormFactors command and transformed to log2 counts per million with a prior count of 1 using cpm command from edgeR package.^45^ PCA was performed on the 2,000 most variable genes after centering and scaling gene expression values across samples. Sample-level clustering was performed using k-means clustering on the first five principal components, with three clusters and 50 random starts. Genes were considered significantly differentially expressed if they had an absolute log2 fold change > 0.5 and an adjusted p value < 0.05.

**Supplemental Figures**
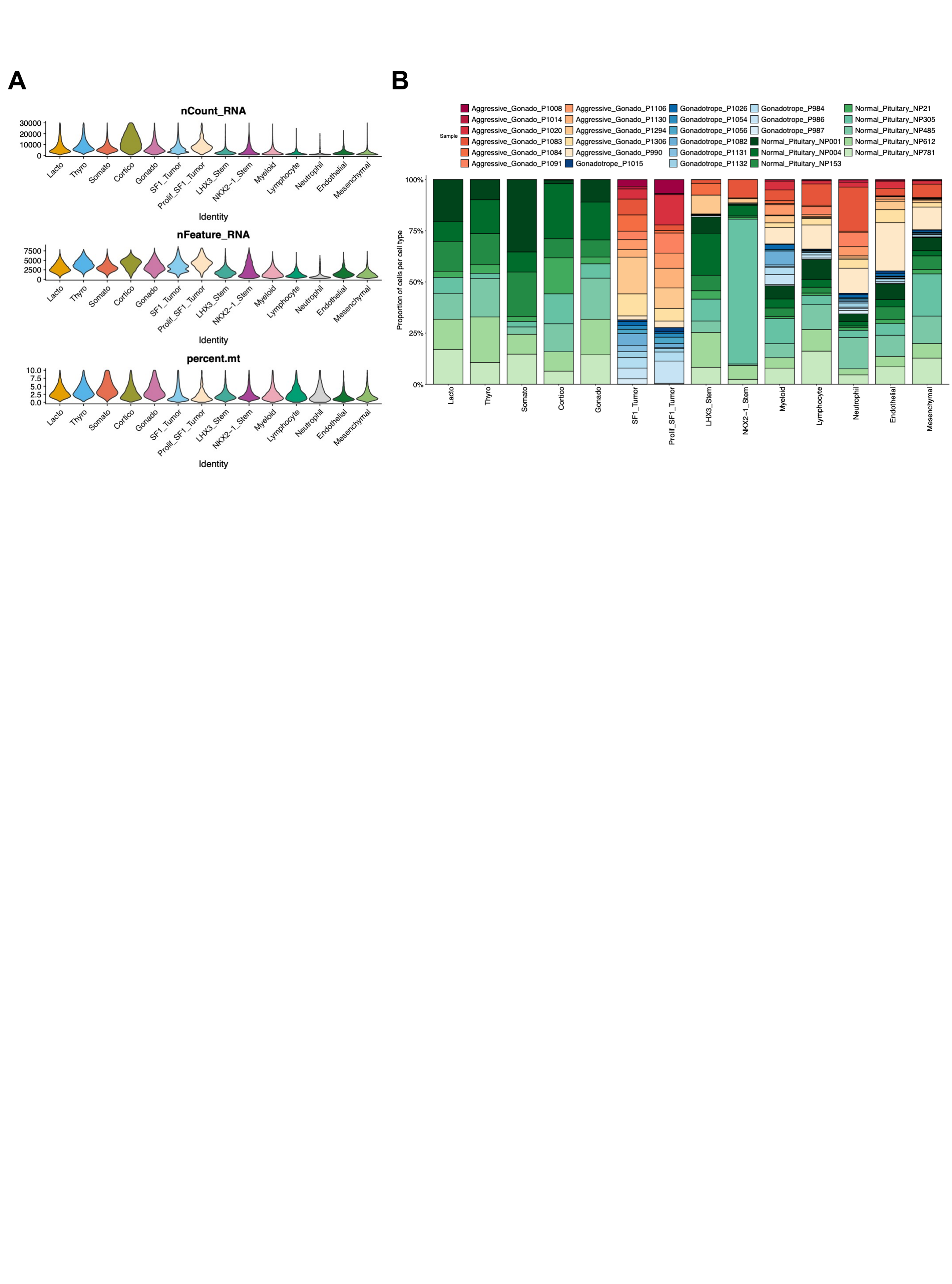

**Supplemental Figure 1. Cell quality metrics and sample composition across cell types in the integrated dataset.** **A.** Violin plots showing cell quality matrix across cell types. nCount_RNA, top; nFeature_RNA, middle; percent.mt, bottom. **B.** Stacked bar plot showing the proportion of cells contributed by each sample within each cell type.

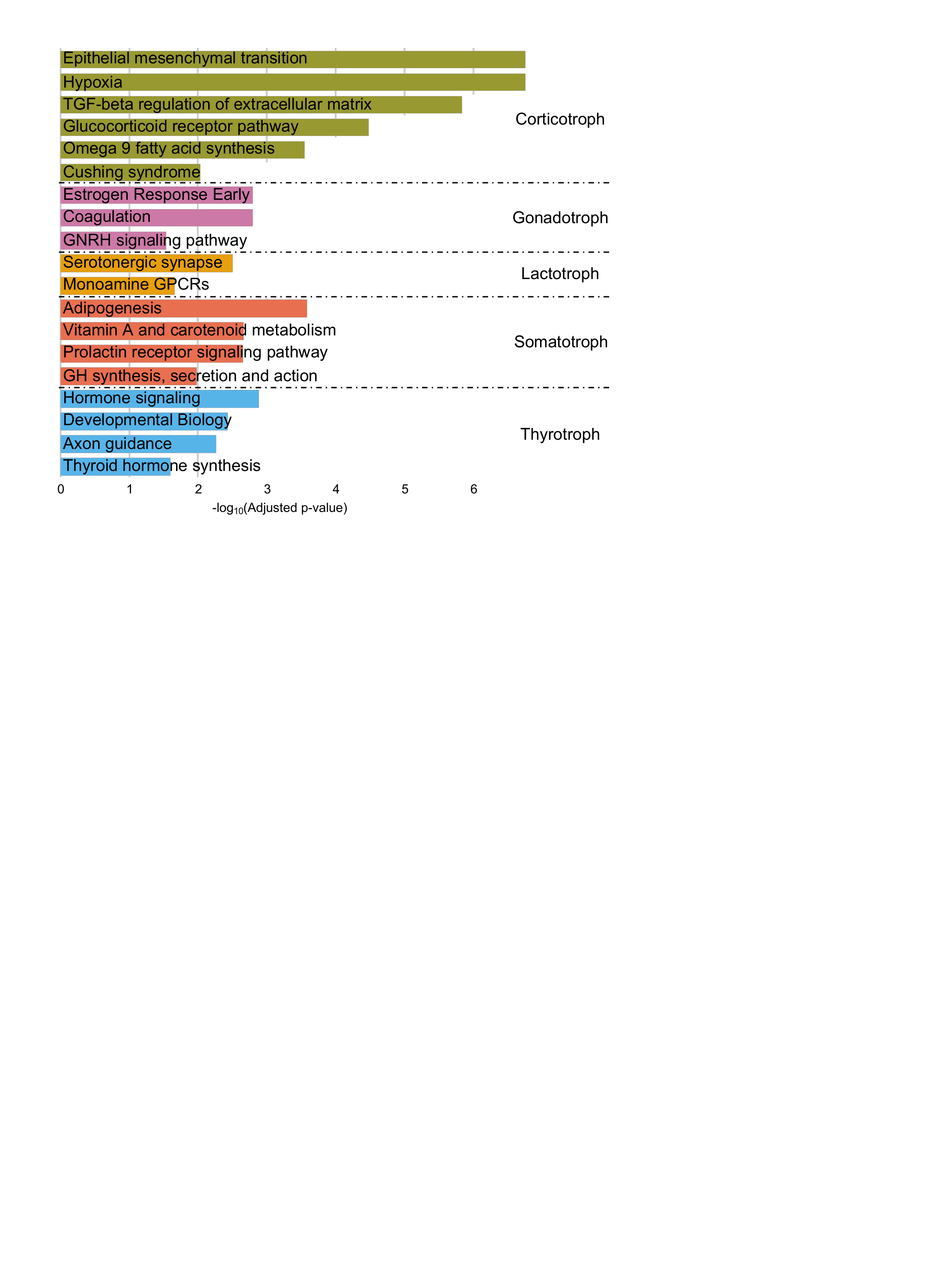

**Supplemental Figure 2. Lineage specific biological pathway enrichment in normal pituitary cells.** Bar plots showing biological pathways enriched in each normal pituitary lineage cell type. Each color represents a distinct pituitary cell lineage

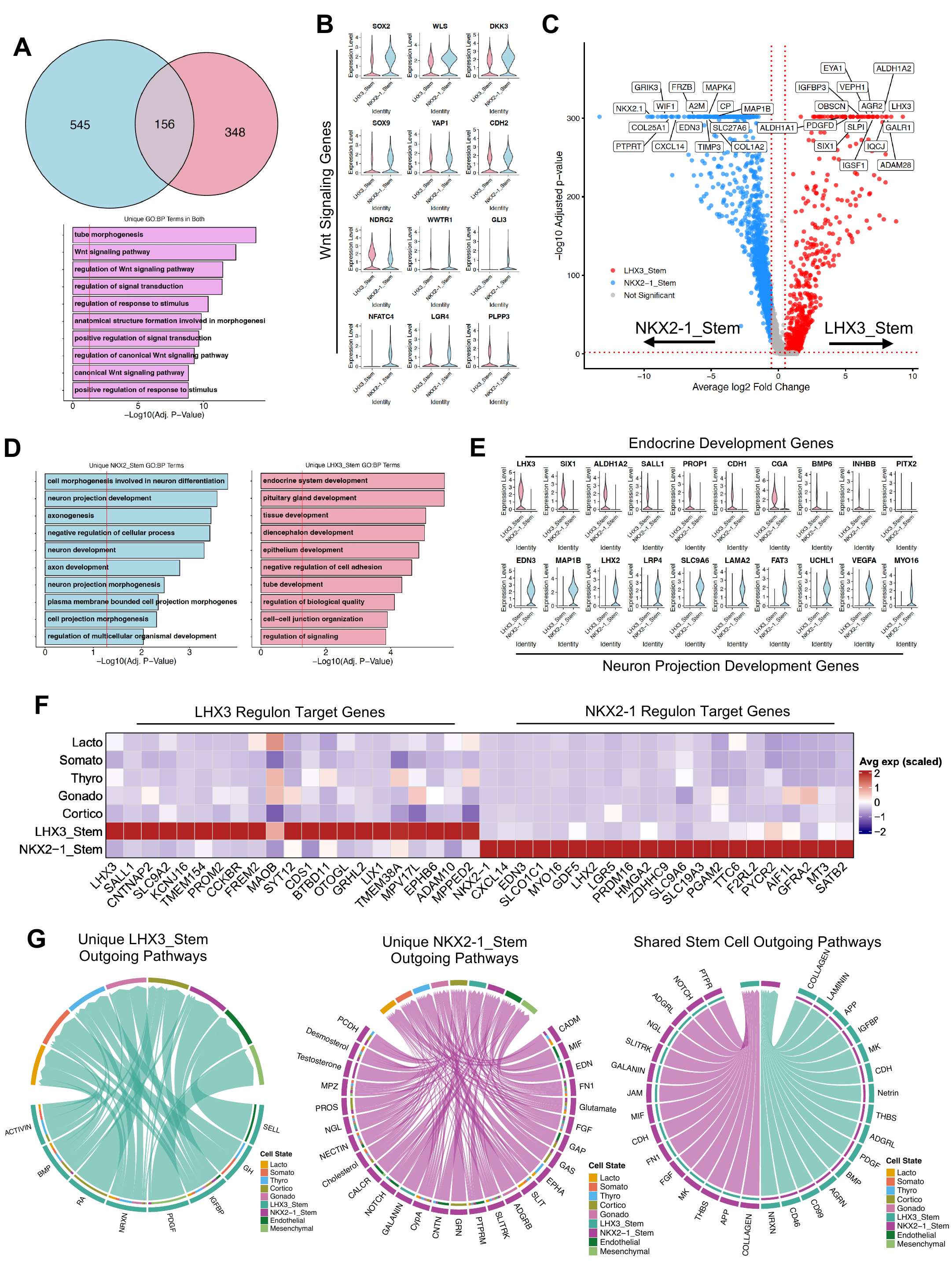

**Supplemental Figure 3. Transcriptomic comparison of stem cell populations in the human pituitary gland.**

**A.** Venn diagram of genes shared between the two stem populations and Gene Ontology (GO) term enrichment of the overlapping genes. **B.** Violin plot showing expression of shared genes driving the common Wnt signaling GO terms. **C.** Volcano plot of differentially expressed genes between the two stem cell populations, *NKX2-1* and *LHX3* stem cells. **D.** Enrichment of top unique GO terms in each stem cell population. **E.** Expression of genes driving the Endocrine Development (upregulated in LHX3 stem cells) and the Neuron Projection Development (upregulated in NKX2-1 stem cells). **F.** Heatmap showing top target genes of the LHX3 and NKX2-1 regulons**. G.** Unique and shared cell-cell communication between the two stem cell populations and other cell types at the aggregate pathway level.

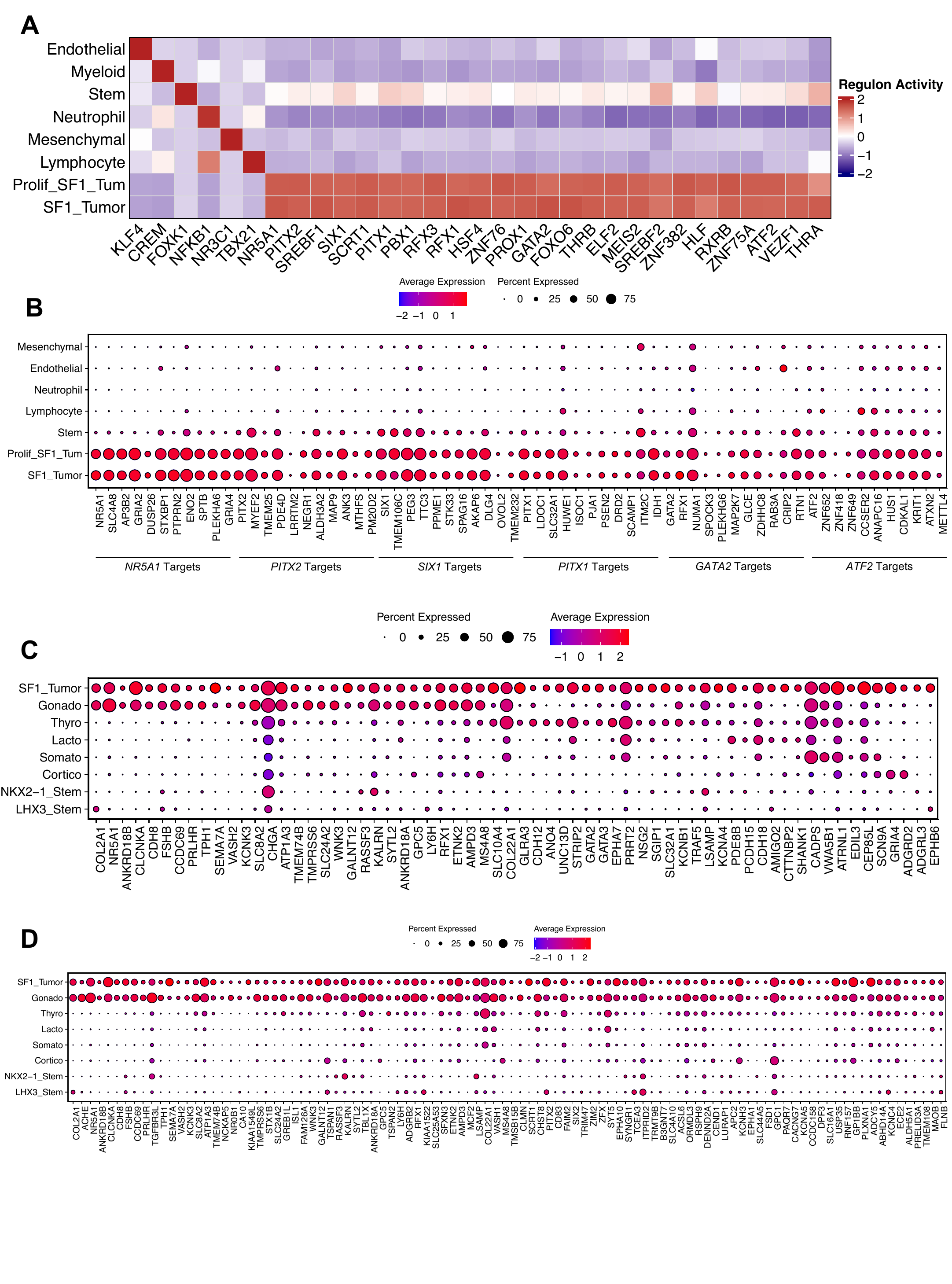

**Supplemental Figure 4. Regulon analysis of non-functioning gonadotroph tumor cells.**
**A.** Heatmap of the top regulon across cell types in the NFG dataset and top 20 regulons in SF1 tumor cells. **B.** Dot plot showing expression of top 10 target genes of the *NR5A1, PITX2, SIX1, PITX1, PROX1,* and *ATF2* regulons across cell types. Dot color and size indicate average expression and the percentage of expressing cells, respectively. **C.** Dot plot showing expression of genes overlapping between SF1 tumor cells and each corresponding normal cell type contributing to the hypergeometric enrichment shown in Fig 3E. **D.** Dot plot showing expression of gonadotroph marker genes identified from the full set of differentially expressed genes in SF1 tumor cells, including genes outside the top 200 used for the hypergeometric enrichment analysis. Only the APG sample sequenced with single-cell RNAseq was used for comparison to tumor samples in C-E.

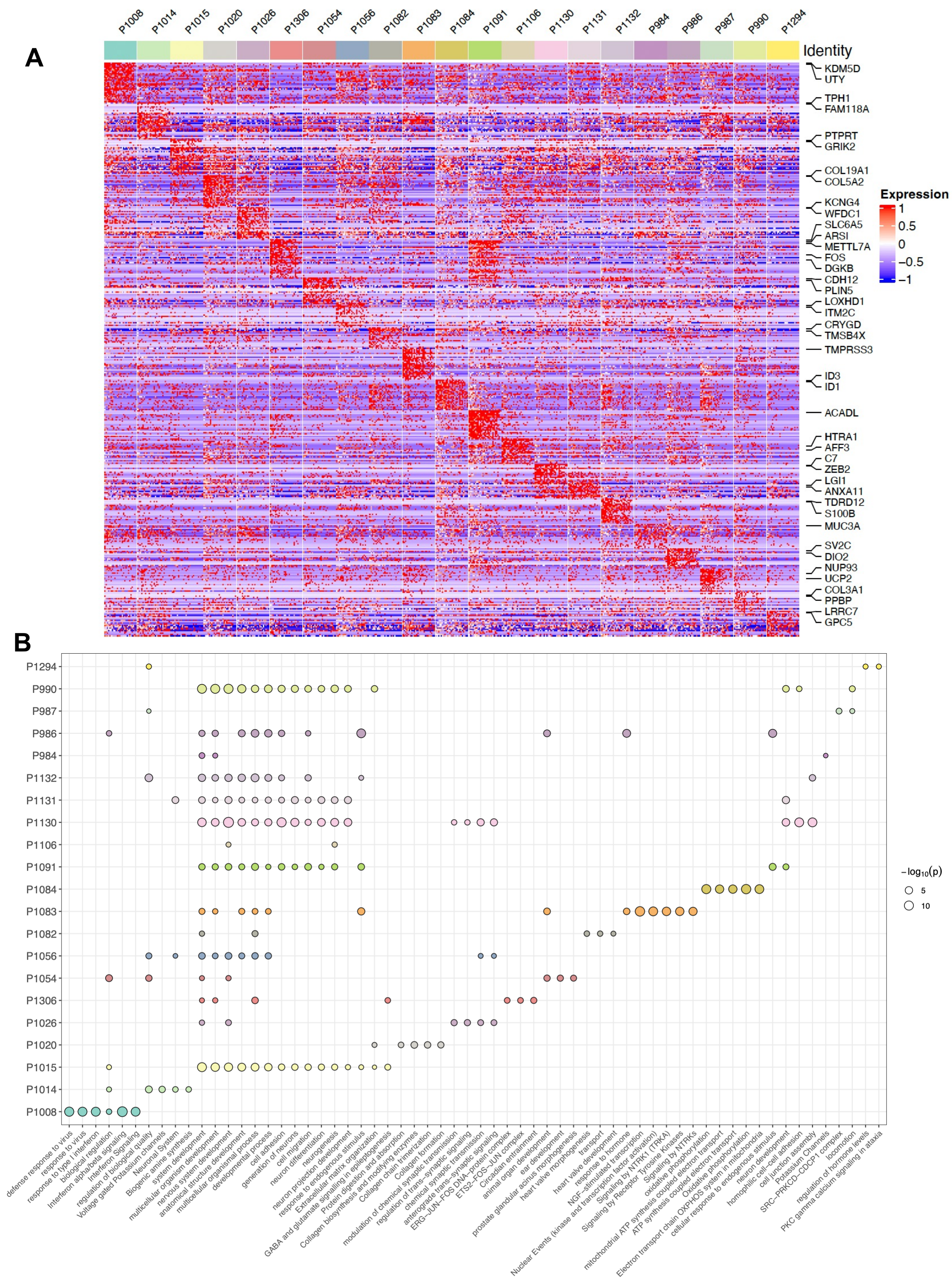

**Supplemental Figure 5. Sample-specific transcriptional programs and pathway enrichment.** **A.** Heatmap showing genes specifically expressed in each tumor sample. **B.** Dot plot showing shared and sample-specific biological pathways across tumor samples.

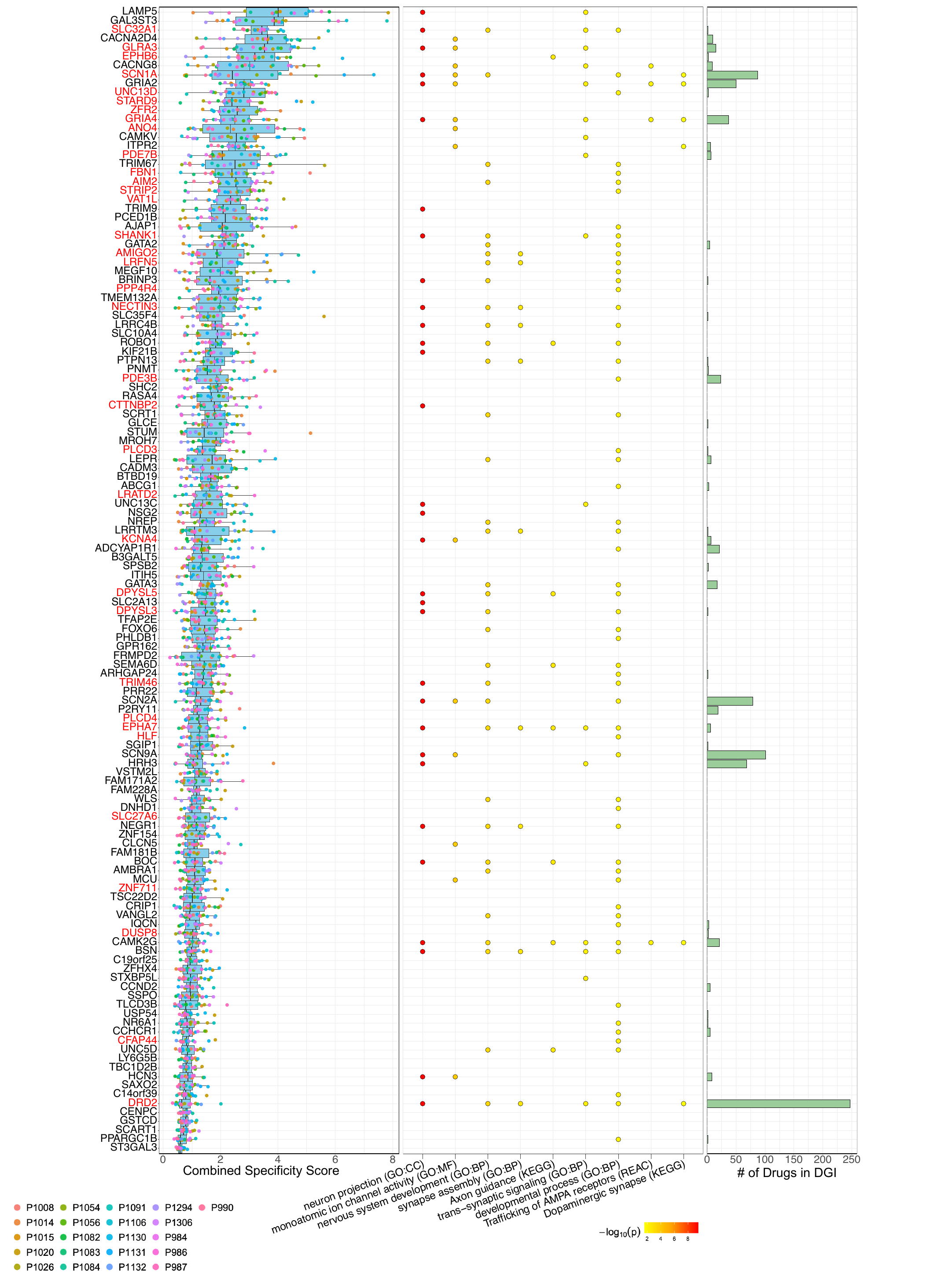

**Supplemental Figure 6. Conserved SF1 Tumor Markers across samples.** (Left) Box plot of combined specificity score of top 100 conserved SF1 tumor markers. Combined specificity score calculated as (percent of tumor cells expressing gene) ∗(1-percent of gonadotroph cells expressing gene)∗avg_log2FC. (Center) Dot plot showing gene inclusion of select ontological terms enriched in among conserved SF1 tumor markers. (Right) Bar plot showing number of drug-gene interactions per gene in the Drug Gene Interaction (DGI) database.

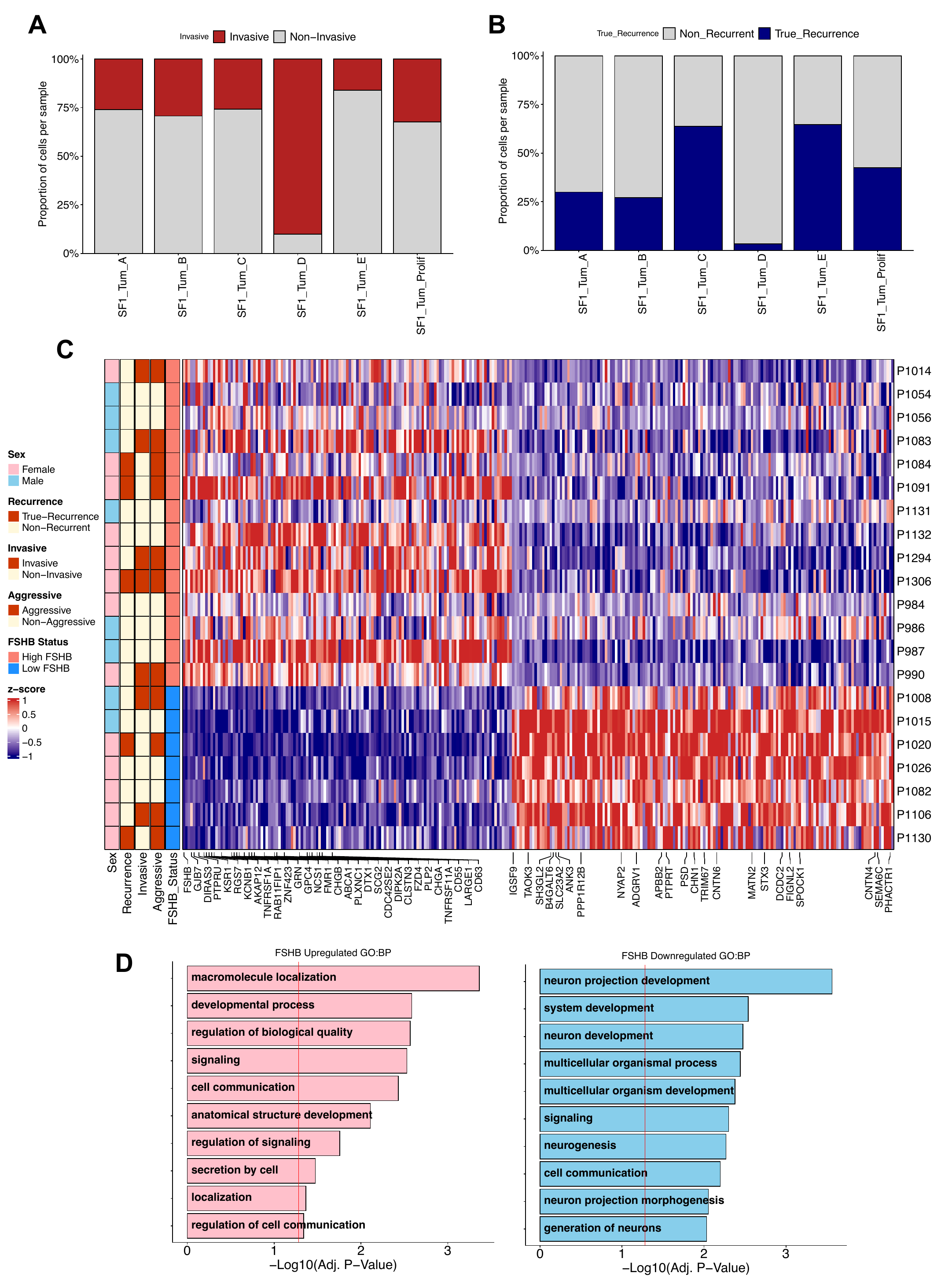

**Supplemental Figure 7. Transcriptional characteristics of clinically defined subgroups. A,B.** Stacked bar plots showing the proportion of cells from invasive versus non-invasive tumors **(A)** and recurrent versus non-recurrent tumors **(B)**. **C.** Heatmap showing scaled normalized pseudobulked counts per sample of all significantly differentially expressed genes identified by pseudobulk analysis between high_FSHB and low_FSHB tumors. Labeled genes are top transcripts driving enrichment of the GO Bioloigical Process terms “signaling” and ”neuron projection development” in high_FSHB and low_FSHB samples respectively. **D.** GO Biological Process enrichment of differentially expressed genes in high_FSHB (left)and low_FSHB (right) samples. Red vertical line indicates significance.

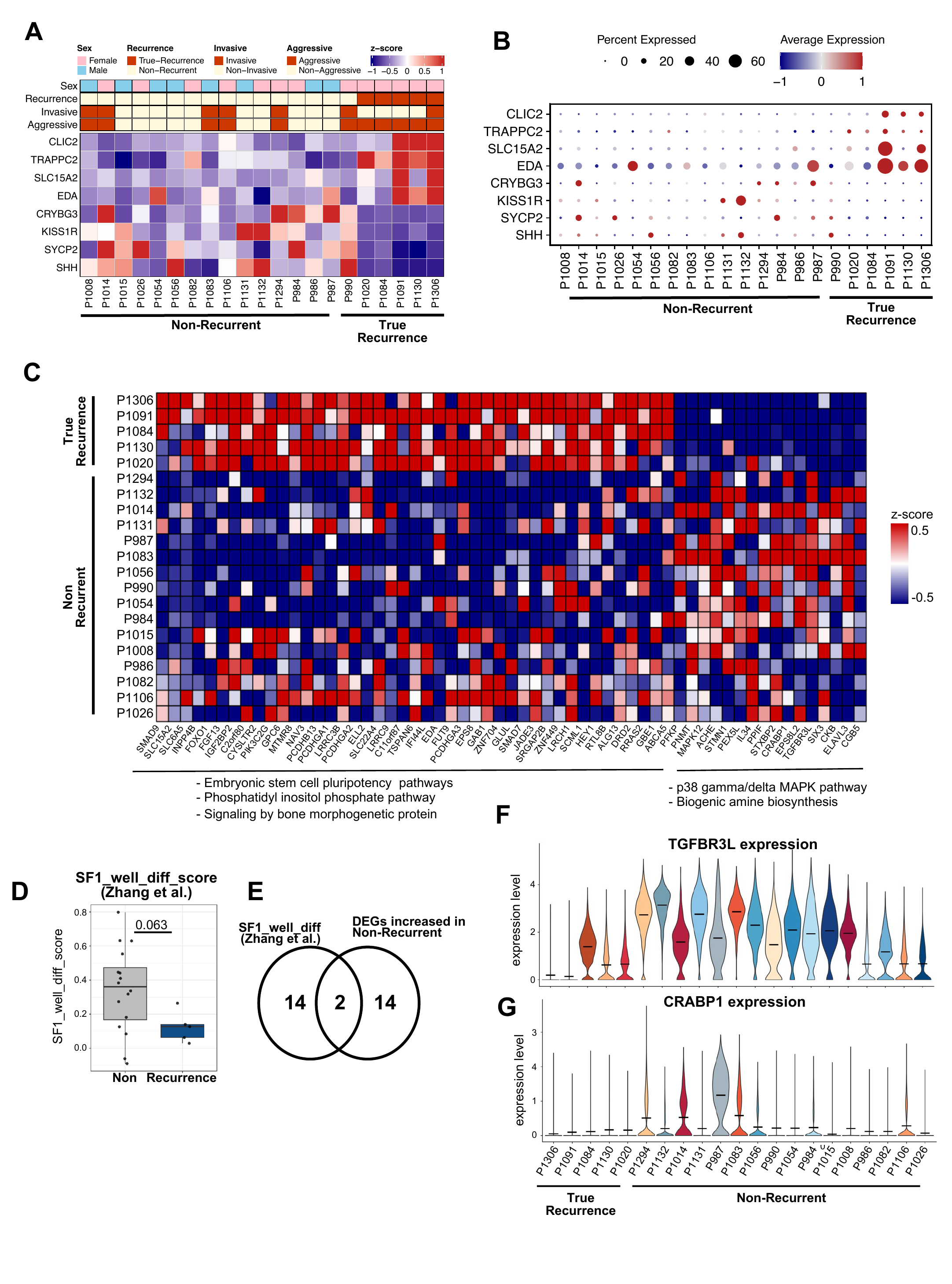

**Supplemental Figure 8. Transcriptional features associated with tumor recurrence.**

**A.** Heatmap showing scaled normalized pseudobulked counts per sample of significantly differentially expressed genes identified by pseudobulk analysis between recurrent and non-recurrent tumors. **B.** Dot plot showing expression of genes pictured in A at the single-cell level. **C.** Heatmap showing sample-level average expression of genes identified as differentially expressed between recurrent and non-recurrent tumors at the single-cell level. Representative pathways enriched in genes upregulated in recurrent (left) and non-recurrent (right) tumors are listed below. **D.** Box plot of the SF1 well-differentiated signature score (Zhang et al.) in non-recurrent and recurrent tumors. Each dot indicates an individual sample. **E.** Venn diagram showing overlap between the SF1 well-differentiated signature genes (Zhang et al.) and genes upregulated in non-recurrent tumors. **F, G.** Violin plots showing *TGFBR3L* and *CRABP1* expression per sample.

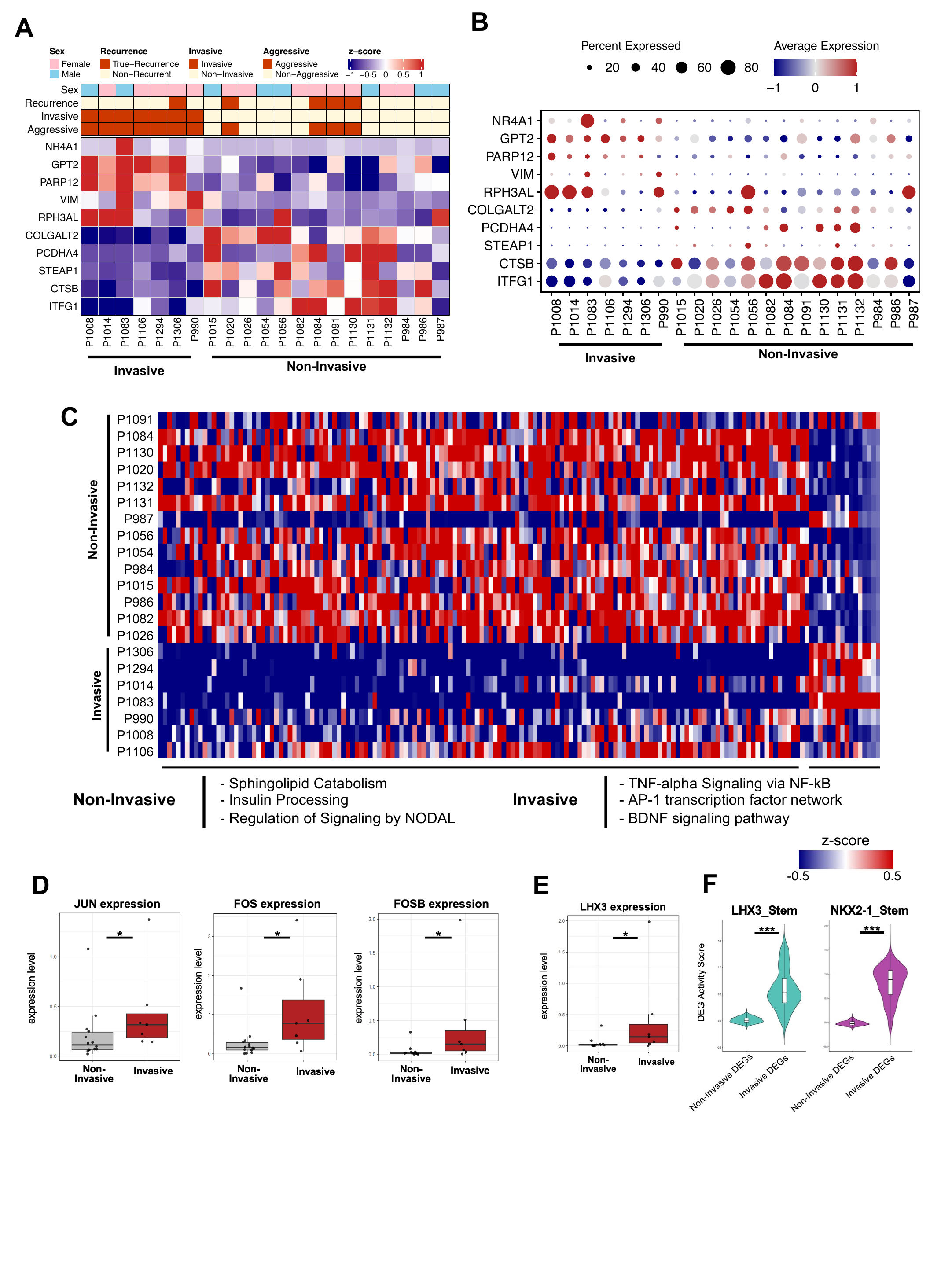

**Supplemental Figure 9. Transcriptional features associated with tumor invasiveness.**

**A.** Heatmap showing scaled normalized pseudobulked counts per sample of significantly differentially expressed genes identified by pseudobulk analysis between invasive and non-invasive tumors. **B.** Dot plot showing expression of genes pictured in A at the single-cell level. **C.** Heatmap showing sample-level average expression of genes identified as differentially expressed between invasive and non-invasive tumors at the single-cell level. Representative pathways enriched in genes upregulated in non-invasive (left) and invasive (right) tumors are listed below. **D.** Box plots showing sample-level average expression of the AP-1 transcription factor genes *JUN* (left), *FOS* (middle), and *FOSB* (right) in non-invasive and invasive tumors. Each dot indicates an individual sample. **E.** Box plot showing sample-level average expression of *LHX3* in non-invasive and invasive tumors. **F.** Violin plots showing signature scores of non-invasive and invasive DEGs in LHX3 stem cells (left) and NKX2-1 stem cells (right). *P < 0.05, ***P < 0.001.

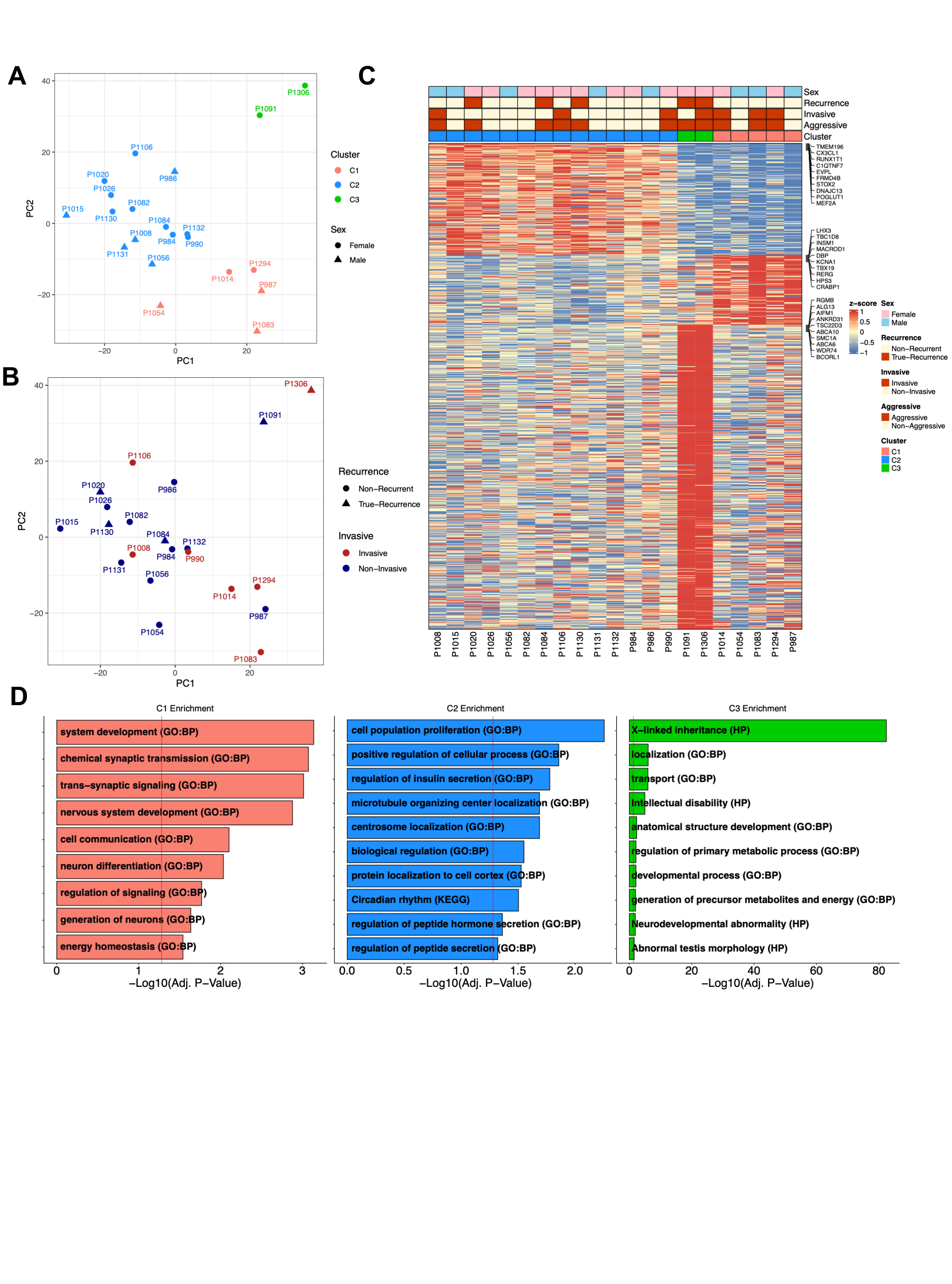

**Supplemental Figure 10. Transcriptome-based classification of non-functioning gonadotroph tumors.** **A, B.** Principal component analysis (PCA) of pseudobulk tumor cell profiles per sample. Samples are colored by k-means cluster (C1-C3) and shaped by sex **(A)**, or colored by invasiveness and shaped by recurrence status **(B)**. **C.** Heatmap showing normalized pseudobulked counts of differentially expressed genes for each cluster per sample. Column annotations indicate sex, recurrence, invasiveness, aggressiveness, and cluster. Representative genes are labeled on the right. **D.** Bar plot showing GO enrichment analysis of genes upregulated in C1 (left), C2 (middle), and C3 (right). GO:BP, Gene Ontology biological process; HP, Human Phenotype Ontology; KEGG, Kyoto Encyclopedia of Genes and Genomes.

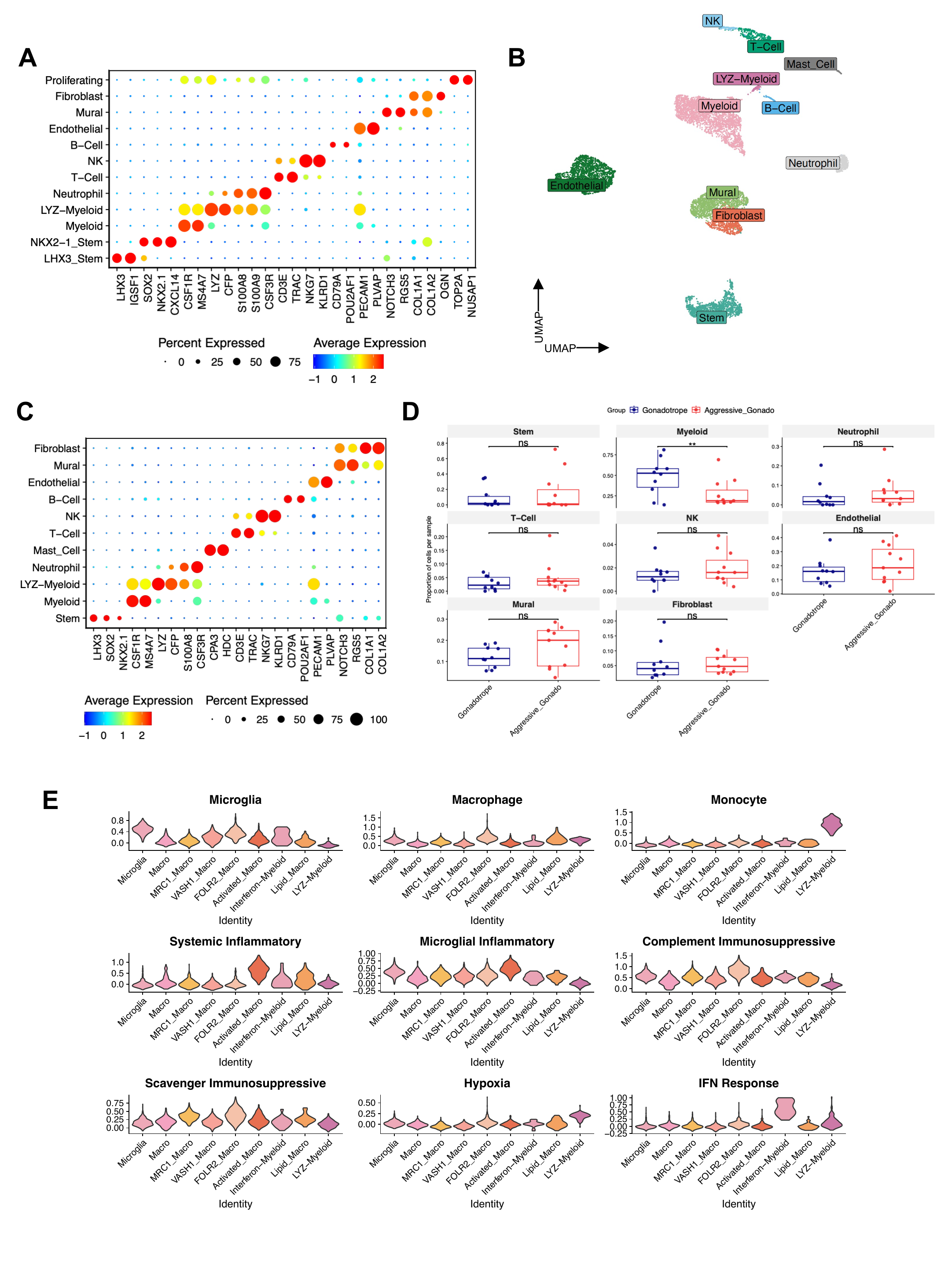

**Supplemental Figure 11. Characterization of tumor microenvironment cell types.** **A.** Dot plot showing marker gene expression across cell types of the integrated NFG and APG microenvironment dataset shown in Fig. 5A. **B.** UMAP of microenvironment cells from NFG samples only. **C.** Dot plot showing marker gene expression across microenvironment cell types from NFG samples. **D.** Box plots showing the proportional differences of each microenvironment cell type per sample in non-aggressive (blue) and aggressive (red) gonadotroph tumors. **E.** Violin plots showing myeloid cell sub-types scored by myeloid consensus programs defined by Miller et al.
